# Microphysiological Drug-Testing Platform for Identifying Responses to Prodrug Treatment in Primary Leukemia

**DOI:** 10.1101/2022.04.06.483760

**Authors:** Furkan Gökçe, Alicia Kaestli, Christian Lohasz, Martina de Geus, Hans-Michael Kaltenbach, Kasper Renggli, Beat Bornhauser, Andreas Hierlemann, Mario Modena

**Affiliations:** ETH Zurich, Department of Biosystems Science and Engineering, Basel, Switzerland; University Children’s Hospital Zurich, Children’s Research Center, Zurich, Switzerland

## Abstract

Despite increasing survival rates of pediatric leukemia patients over the past decades, the outcome of some leukemia subtypes has remained dismal. Drug sensitivity and resistance testing on patient-derived leukemia samples provide important information to tailor treatments for high-risk patients. However, currently used well-based drug screening platforms have imitations in predicting the effects of prodrugs, a class of therapeutics that require metabolic activation to become effective. To address this issue, we developed a microphysiological drug-testing platform that enables co-culturing of patient-derived leukemia cells, human bone marrow mesenchymal stromal cells, and human liver microtissues within the same microfluidic platform. This platform also enables to control the physical interaction between the diverse cell types. We were able to recapitulate hepatic prodrug activation of ifosfamide in our platform, which is very difficult in traditional well-based assays. By testing the susceptibility of primary patient-derived leukemia samples to the prodrug ifosfamide, we identified sample-specific sensitivities to ifosfamide in primary leukemia samples. We found that our microfluidic platform enabled the recapitulation of physiologically relevant conditions and the testing of prodrugs including short-lived and unstable metabolites. The platform holds great potential for clinical translation and precision chemotherapy selection.

## 1. Introduction

Since the initial efforts in the 1940s to 1960s, drug screening for leukemia has heavily relied on high-throughput testing of compound libraries on primary or patient-derived samples, mostly using ex vivo well plate-based assays ^1–3^. The prediction of drug efficacy through such tests has led to a massive decrease in mortality of acute lymphoblastic leukemia (ALL) ^3^. While cell lines established from primary tumors enabled robust and reproducible protocols for drug development and screening studies, they have been limited to recapitulating specific genetic alterations and have been inadequate in representing the genomic diversity and heterogeneity of primary leukemia cases ^4,5^. In addition, some cell lines have been reported to be resistant to anticancer molecules ^6^, which has further reduced their applicability for drug testing. More recently, the growing clinical interest in genomic profiling of primary leukemia cases has greatly helped to understand the lesions that occur in cancer, especially for pediatric ALL, with implications on treatment identification ^7,8^. This improved understanding has led to several functional approaches to develop precision and personalized medicine tools by sequencing and testing primary leukemia cells at single-cell resolution to identify drug response profiles of patients ^9–13^.

Today, the identification of effective therapeutic approaches for high-risk ALL subtypes is rendered challenging by the lack of representative cell-line models and by the relatively low number of patients having the disease ^12^. Disease subtypes are characterized by a variable genetic landscape that is directly affecting the patient responses to treatment, and primary ALL cells can be used to recapitulate the state of the disease of the respective patients in vitro ^8,12,13^. Patient-derived xenografts (PDX) have been widely used as in-vivo models to study the biology of leukemia and have been key in leukemia drug-screening studies, as these models feature characteristics that are very similar to those of primary ALL cells. Using patient-derived xenografts, it is possible to generate biobanks for high-throughput testing of patient-derived cells ^8,12–16^. The PDX models are generated by collecting leukemia cells from patients via liquid biopsy and injecting the collected primary leukemia cells in immunodeficient mice so that the leukemia cells can proliferate and expand, as illustrated in **Figure 1**.a ^14^. The leukemia cells from PDX models (PDX cells) can then be cultured with human bone-marrow-derived mesenchymal cells (MSCs) in serum-free culture medium in well plates and be exposed to anticancer drugs ^8,12^ (**Figure 1**.b). MSCs enhance the in vitro viability of PDX cells by secreting essential signaling and growth factors and have been shown not to alter the obtained results with tested compounds ^17–19^.

**Figure 1.**
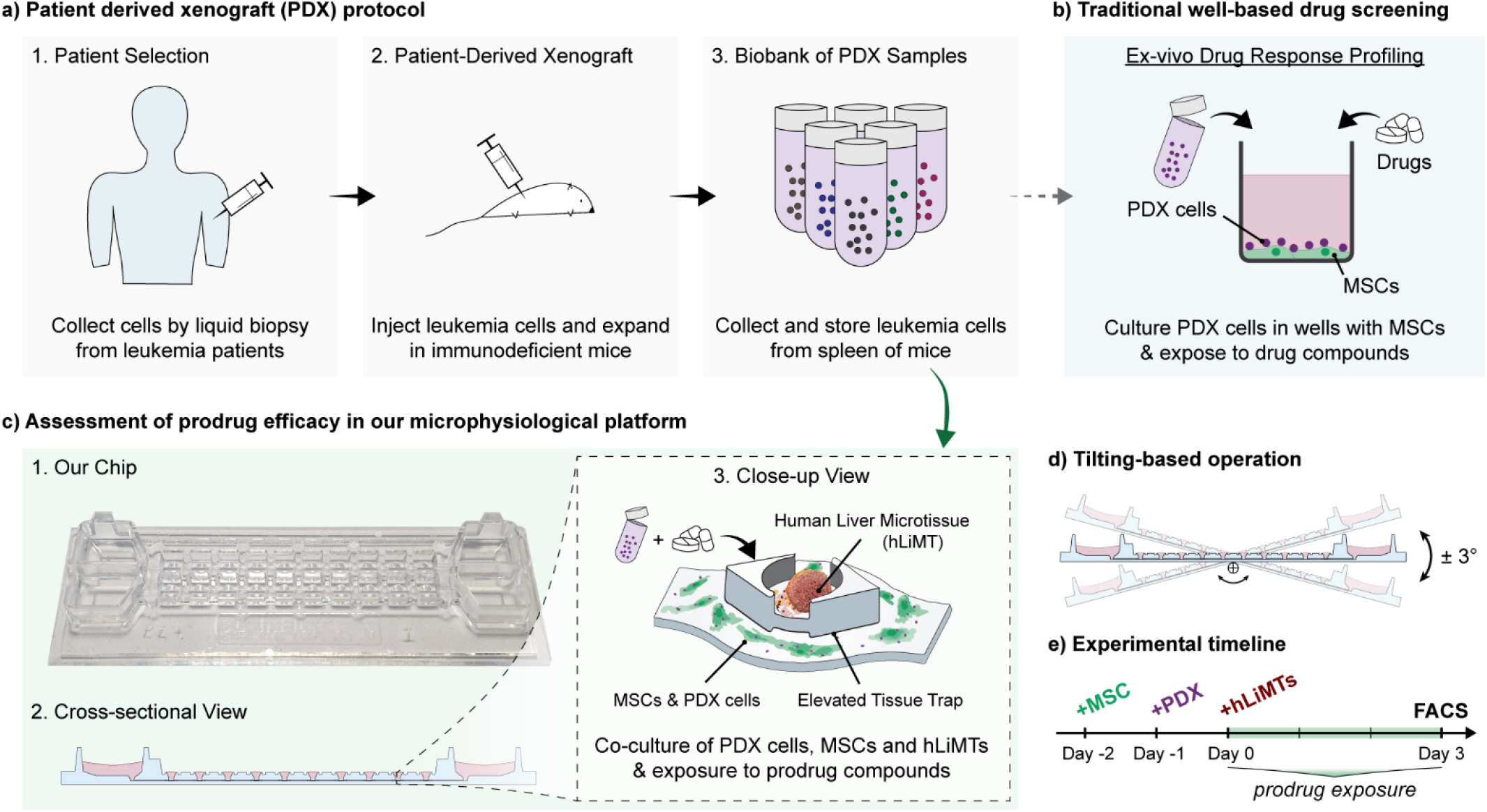
Overview of drug screening experiments using patient-derived xenograft models. **a)** Summary of the protocol for generating patient-derived xenografts: (1) Selection of patients from high-risk acute lymphoblastic leukemia cases and collection of leukemia cells by liquid biopsy. (2) Injection and expansion of collected leukemia cells in immunodeficient NSG mice. (3) Collection and storage of expanded patient-derived leukemia cells from the spleen of xenografted mice and establishment of a biobank of patient-derived xenograft leukemia (PDX) samples. **b)** Traditional well-based drug screening, during which the PDX cells from the biobank are cultured in well plates together with human bone-marrow mesenchymal stromal cells (MSCs) and are then exposed to anticancer drugs. **c)** Assessment of prodrug efficacies in the microphysiological testing platform: (1) A photograph of the platform and (2) a cross-sectional view showing the medium reservoirs at each end, interconnected through a microfluidic channel hosting 10 microtissue compartments. (3) Close-up view of one microtissue compartment hosting a human liver microtissue (hLiMT) in an elevated compartment for the metabolic transformation of prodrugs as well as MSCs and PDX cells at the bottom of the microfluidic channel. **d)** Tilting-based operation of the platform for perfusion, i.e., supply of nutrients and transport of metabolites through gravity-driven flow. **e)** Experimental timeline showing the addition of the distinct cell types to the microfluidic chip, prodrug exposure, and flow-cytometry (FACS) analysis at the end of the experiment after three days of prodrug treatment.

Well plate-based, high-throughput assays are widely used today in functional precision medicine to test antileukemia efficacies of drug compounds, and robotic handling of assays has allowed for increasing the testing throughput ^8,10,12^. However, despite the increased throughput, the overall testing strategy has remained largely unchanged since the 1960s. In these tests, only direct anticancer effects of the compounds are tested, an approach that falls short of recapitulating key physiological phenomena in the body, such as drug metabolic processes that affect drug action and efficacy ^20,21^.

Upon administration to a patient, a drug interacts with many tissue types, through <u>a</u>bsorption (e.g., in the gut, lung, skin), <u>d</u>istribution (e.g., by the blood or interstitial flow), <u>m</u>etabolism (e.g., by enzymes in the liver) and <u>e</u>xcretion (e.g., by the kidney, lung), the four key processes of pharmacokinetics typically summarized by the acronym ADME ^20,22^. For a particular class of drugs, the so-called prodrugs, enzymatic or chemical reactions are required to convert the compounds into their active form ^23^. Prodrugs have accounted for over 10% of newly approved therapeutic molecules in the last decade owing to, e.g., their favorable water solubility, improved stability, longer half-life, or better passive permeability ^23^. Several prodrugs, such as oxazaphosphorines, have already been proven highly efficacious for leukemia treatment ^24^. Many of those prodrugs require metabolic activation by cytochrome P450 (CYP) enzymes that are highly expressed in the liver ^23,24^. CYP enzymes are also necessary for the activation and inactivation of both, anticancer drugs and supportive substances or adjuvants ^25^. For instance, epipodophyllotoxins and ifosfamide are anticancer drugs that require CYP enzymes for activation, whereas glucocorticoids and vincristine are anticancer drugs that require CYP enzymes to be inactivated ^25^. Some prodrugs have stable metabolites, such as irinotecan, and therefore can easily be tested in vitro by exposing the target cells directly to their metabolites, such as SN-38, the active metabolite of irinotecan ^26^. However, some other prodrugs feature unstable or short-lived metabolites, such as nitrogen mustards in the case of oxazaphosphorines, that render in vitro testing particularly challenging ^27^. It is, therefore, vital to include prodrugs and their metabolic activation in drug screening studies to effectively assess the impact of short-lived metabolites on leukemia cells.

Microphysiological systems (MPSs), also known as organ-on-chip devices, are microfluidic multi-tissue culture platforms that recapitulate the physiology of parts of the human body and organ-specific functions in vitro. They include living cell and tissue cultures and enable cell-cell and tissue-tissue interactions through microfluidic coupling ^20,21^. By recapitulating complex physiological functions and tissue-tissue interactions of the human body in vitro, MPSs constitute more representative preclinical models of ADME for pharmacology and toxicology studies. MPSs offer the potential of improving disease modeling and conducting personalized-medicine studies in vitro, as they help to avoid issues related to interspecies differences resulting from the use of animal models ^20,21,28–30^. In particular, liver-on-chip MPSs, which mimic the physiological structure and function of liver, have been demonstrated to be more predictive models for pharmacology and toxicity studies in comparison to their 2D, well-based counterparts ^30–32^. 3D liver-tissue models more accurately recapitulate the structure and function of the organ in vitro and can be integrated in microfluidic networks and combined with other tissues. The resulting MPSs feature rapid delivery of nutrients and exchange of signaling factors ^32–35^. For instance, a multiorgan MPS was developed to recapitulate the metabolism of tolcapone, a drug used to treat Parkinson’s disease ^36^. The MPS enabled the identification of twelve tolcapone metabolites and key biomarkers in the brain, including three metabolites that have not been reported until then. MPSs also hold the promise to promote developments in personalized medicine by offering the possibility to co-culture primary and patient-derived samples with relevant tissue or organ models in vitro for studying, e.g., tumor microenvironments to identify key biomarkers or for profiling drug responses of individual patients to select treatment options ^28,37,38^. Although several MPSs have been reported for studying toxicity and metabolism of drugs and nanomedicines ^29–31,39–41^, there are only a few that feature metabolic activation of prodrugs for screening the effects of drug metabolism on diseased target cells in a single self-contained system ^33,42–46^.

In this study, we present a microphysiological drug-testing platform featuring the capability to test liver-mediated prodrug transformation on patient-derived leukemia samples. To ensure effective testing with the intermediary and final-stage metabolites of a prodrug producing short-lived metabolites, the leukemia cells need to be cultured in close vicinity to the metabolically active tissue, i.e., liver microtissues. Nevertheless, the leukemia cells and liver microtissues must be kept physically separated during the drug exposure study to avoid confounding effects of direct physical interaction between the different cell types. To this end, we realized a multi-tissue microfluidic platform that enables the co-culturing of leukemia cells and liver microtissues relying on tubing- and pump-free medium recirculation. Our platform improves on existing drug screening technology by (1) enabling co-cultures of leukemia cells and liver microtissues under conditions of minimized direct physical interaction in the same system to recapitulate hepatic drug metabolism and by (2) enabling continuous medium perfusion in the system through gravity-driven flow. The latter is necessary to establish stable and reproducible conditions and to avoid potential diffusion-limitation in the transport of signaling molecules and drug metabolites. To the best of our knowledge, this is the first MPS platform that offers the possibility to activate prodrugs that yield short-lived metabolites for the treatment of primary leukemia samples and for profiling of their responses ex vivo in a single system. We first characterized our platform and then executed proof-of-concept experiments to test the susceptibility of primary leukemia cells to the prodrug ifosfamide. We used patient-derived xenograft (PDX) models of standard- and high-risk cytogenetic subtypes of pediatric acute lymphoblastic leukemia (ALL) as test samples. We screened multiple patient-derived samples and found that the efficacy of ifosfamide through hepatic activation by the human liver microtissues (hLiMTs) and the production of short-lived metabolites could be assessed in our platform. Moreover, by using our platform, we identified one patient sample that showed a stronger response to ifosfamide treatment with hLiMTs in co-culture. We confirmed the specific sample response to ifosfamide metabolites by co-dosing ritonavir, a known CYP3A4 inhibitor. A corresponding specific response to ifosfamide could not be found with the well-based screening assay.

## 2. Results

### 2.1. Establishing the platform for ex vivo testing of prodrugs

We developed a platform with medium reservoirs at each end of a microfluidic channel hosting 10 microtissue compartments (**Figure 1**.c, **Supplementary Figure 1**). Four of these microtissue compartments represented a metabolic unit and each of them harbored a human liver microtissue (hLiMT) (**Supplementary Figure 2**), while the microfluidic channel was used to culture PDX cells and the MSC feeder layer (**Figure 1**.c). Each chip contained two parallel microfluidic networks and featured standard microscopy slide dimensions for straightforward integration with existing microscopes and chip holders (**Figure 2**). The platform design was developed on the base of a polystyrene-based microfluidic chip devised previously by our group ^22^. In the new design, the microtissue compartments were elevated by 20 μm with respect to the interconnecting microfluidic channel so as to physically separate co-cultured microtissues and adherent cells and to hinder PDX cells and MSCs from entering the tissue compartment (**Figure 1**.c, **Supplementary Figure 1**, **Supplementary Figure 2**). Tissue-tissue or tissue-cell interaction was possible through the common liquid phase and continuous liquid perfusion.

**Figure 2.**
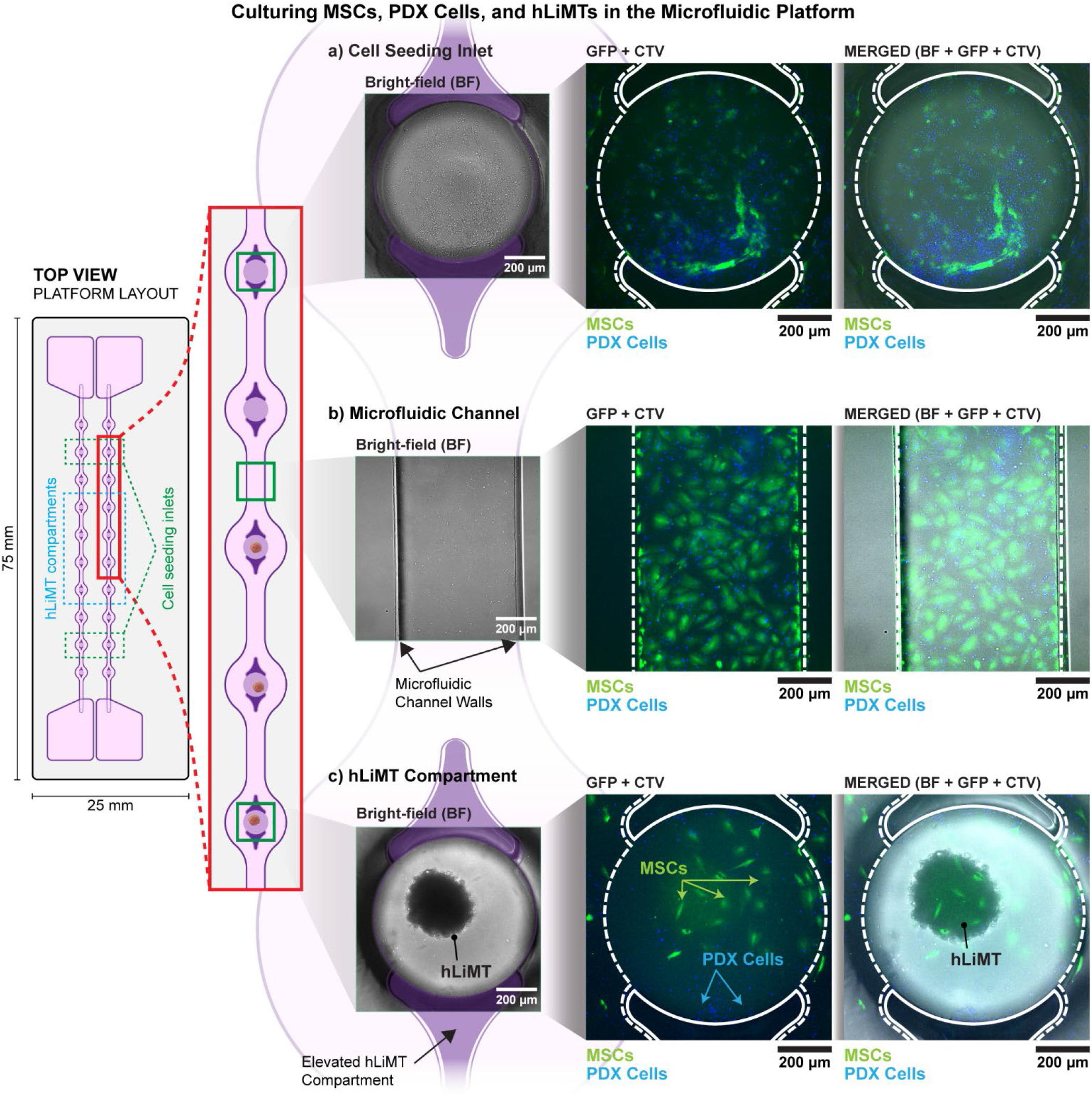
Microfluidic layout of the platform, including a top view of the platform and a close-up view of one microfluidic culture chamber. A platform features two parallel microfluidic culture chambers on a standard-microscopy-slide-size chip. Microscopy images show MSCs, PDX cells, and a hLiMT in the platform (in different regions of the culture chamber, which are indicated by green square boxes in the close-up view) three days after seeding MSCs. <u>LEFT</u>: Bright-field (BF) images, overlayed with the channel layout to show the walls of the microfluidic channel and the elevated hLiMT platform, <u>CENTER</u>: GFP+CTV channels merged to show GFP-expressing MSCs, and PDX cells stained with CellTrace Violet (CTV), <u>RIGHT</u>: overlays of all channel images (BF+GFP+CTV). **a)** MSCs (green) and PDX cells (blue) in the compartment near the seeding inlet). **b)** MSCs (green) and PDX cells (blue) at the bottom of the interconnecting microfluidic channel, showing PDX cells in direct physical interaction with the MSCs. **c)** A hLiMT in the elevated compartment. The elevated platform minimized direct physical interaction between hLiMTs and MSCs/PDX cells by preventing MSCs/PDX cells from growing near the hLiMTs. The elevated hLiMT compartment and channel walls are indicated in the microscopy images.

The platform was operated on a tilting device, which generated a gravity-driven flow due to height differences between the two media reservoirs at the ends of the microfluidic channels. The hydraulic pressure difference between the reservoirs induced a pump- and tubing-free perfusion of nutrients and metabolites inside the channels ^22^ (**Figure 1**.d). Parallelized experimentation was enabled by stacking multiple platforms and simultaneous operation on a single tilting device (**Supplementary Figure 3**).

To perform drug testing including metabolic activity with PDX cells in our platform, we adapted established protocols for well-based drug profiling experiments ^8,12^. Specifically, we maintained the same MSC-to-PDX cell ratio (1:10), while the absolute number of seeded cells was scaled to compensate for the variation in active surfaces for cell attachment. The active surface in our platform was approximately 4 times larger than that in the originally employed well of a 384-well plate. For the control experiments, we opted for using a 96-well plate, as then each well featured the same surface area as our microfluidic platform. This choice also enabled to use the same medium volume, 200 μL, for both the microfluidic platform and the well-plate control.

Drug testing experiments were carried out over 5 days (**Figure 1**.e): MSCs and PDX cells were first seeded in the microfluidic channel during the first two days (Days −1 and −2). On Day 0, the hLiMTs were loaded into the tissue compartments. The prodrug was then added to the system, and all cells were exposed to the test compounds and their metabolites for 3 days. PDX cell viability was assessed at the end of the assay by collecting the cells from the channels and from the wells, and by using flow-cytometry analysis.

Microscopy imaging of the chips three days after seeding of the MSCs confirmed that PDX cells and MSCs adhered to the compartments connected to the cell-seeding inlets (**Figure 2**.a) and the microchannel surface (**Figure 2**.b). MSCs and PDX cells were in direct physical contact, mimicking the standard in-well culture conditions aimed at supporting PDX cell viability. In contrast, very few cells were present in the metabolic compartments, into which the hLiMTs were loaded (**Figure 2**.c). The images validated the function of the elevated stage in the compartment, which physically separated hLiMTs from the MSC/PDX co-culture so that direct physical interactions between the cell and tissue models were minimized during the assay.

### 2.2. Characterization of liver metabolism and bioactivation of ifosfamide

To recapitulate hepatic drug metabolism in vitro, we used commercially available human liver microtissues (hLiMTs, InSphero AG, Schlieren, Switzerland) formed from primary hepatocytes. As a test compound, we used the prodrug ifosfamide, an oxazophosphorine, which was specifically designed to require cytochrome P450 (CYP) activation for the generation of highly active but short-lived metabolites that attack DNA molecules in target cancer cells ^24,27^. Ifosfamide is metabolized mainly by the CYP3A4 and somewhat by the CYP2B6 enzymes to 4-hydroxyifosfamide (IF-OH), which, after a cascade of biochemical reactions, finally decays inside cancer cells to the toxic isophospharamide mustard (iso-PM) (**Figure 3**.a) ^24,47,48^.

**Figure 3.**
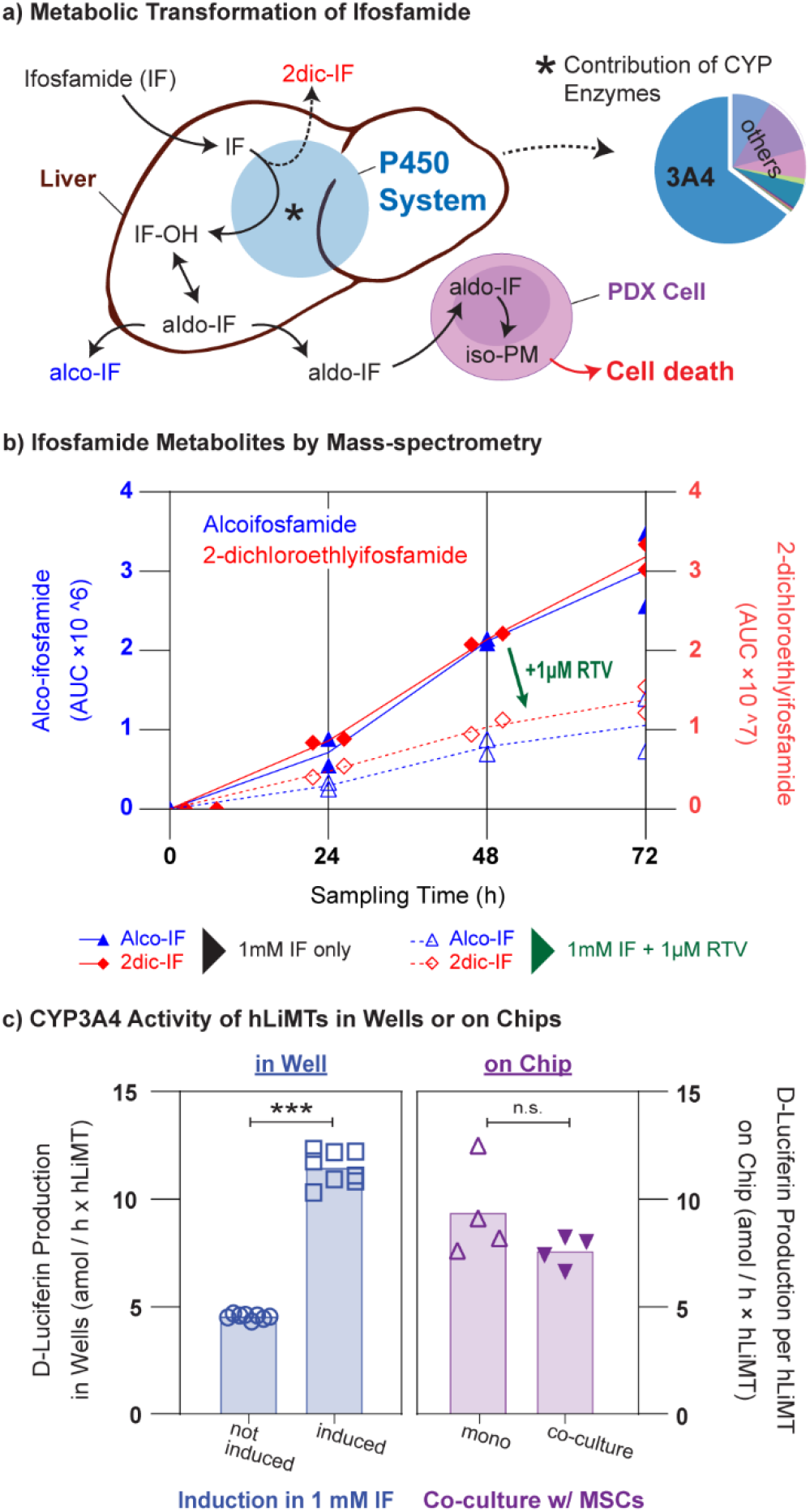
Characterization of the liver metabolism for prodrug transformation. **a)** Metabolic pathway of ifosfamide transformation by the cytochrome P450 system in the liver and contribution of CYP3A4 in comparison to other CYP enzymes. Ifosfamide is first hydroxylated by the P450 system to 4-hydroxyifosfamide (IF-OH) that is in equilibrium with its tautomer aldo-ifosfamide (aldo-IF), which spontaneously decays inside leukemia cells to the active form of interest, isophospharamide mustard (iso-PM) a DNA-crosslinking agent, which is toxic for fast-dividing cells. **b)** Quantification of ifosfamide metabolites generated by hLiMTs in culture well plates by mass spectrometry with or without inhibition of CYP3A4 activity by ritonavir (n=2 replicates in wells). Ritonavir drastically slowed down the ifosfamide metabolism. Concentrations were monitored over 72 h at intervals of 24 h. **c)** Measured CYP3A4 activity in a well plate and in our platform using the P450-Glo CYP3A4 assay. <u>LEFT</u>: D-luciferin production over 72 h by hLiMTs maintained in individual wells, with and without 24-h induction with 1 mM ifosfamide before the CYP3A4 assay (n=8 replicates in wells). <u>RIGHT</u>: D-luciferin production per hour × hLiMT over 72 h by four hLiMTs, maintained in individual microfluidic culture chambers either in monoculture or in co-culture with 10k MSCs in our platform (n=4 replicates on chip). All hLiMTs for the on-chip measurements were induced for 24 h in 1 mM ifosfamide solution before the CYP3A4 assay. D-luciferin production was normalized to the number of MTs in order to compare the average CYP3A4 activity in wells and on chips. *** P<0.001, n.s. = not significant. The samples were compared using a two-tailed unpaired t-test (Welch’s t-test).

To validate the hLiMT ability to metabolize ifosfamide, we exposed the hLiMTs in well plates to 1 mM ifosfamide and measured the production of ifosfamide metabolites over 72 h by using mass spectrometry (MS). Our results confirmed the production of several metabolites of ifosfamide via liver-mediated transformation including alco-ifosfamide and 2-dichloroethylifosfamide (**Figure 3**.b). Alco-ifosfamide is a metabolite of the drug-activation pathway of ifosfamide, whereas 2-dichloroethylifosfamide is a metabolite of the drug-deactivation pathway ^24,48^. Production of both metabolites confirmed and validated the capability of hLiMTs to recapitulate liver-mediated prodrug transformation in vitro. The metabolite concentrations showed a linear increase throughout 72 hours. To confirm that metabolites were a product of liver transformation, we also exposed the hLiMTs to both ifosfamide and ritonavir. Ritonavir is a known CYP3A4 inhibitor and reported to inactivate the CYP3A4 enzymes by irreversibly binding to the active site of the enzyme ^49^. The addition of ritonavir to the medium resulted in three-fold lower concentrations of ifosfamide metabolites after 72 h in comparison to wells with ifosfamide only, which confirmed the role of the CYP3A4 enzyme in the conversion of ifosfamide (**Figure 3**.b).

Next, we characterized the CYP3A4 activity in standard well plates and on-chip by measuring the average conversion of Luciferin-IPA to D-luciferin per hour per hLiMT. We first investigated the effect of inducing the metabolic activity of hLiMTs by administering 1 mM ifosfamide. We observed that a 24-h treatment of hLiMTs with the drug solution resulted in a more than 2-fold average CYP3A4 activity during 72 h (**Figure 3**.c), from 4.54 ± 0.12 amol/h D-luciferin for non-induced tissues to 11.46 ± 0.77 amol/h for induced ones (P<0.001). We then cultured previously induced hLiMTs on-chip (4 hLiMTs per channel) for 3 days either in monoculture or in co-culture with 10k MSCs. Our measurements showed no significant difference in CYP3A4 activity for the two conditions (9.35 ± 2.19 amol/h per hLiMT and 7.56 ± 0.72 amol/h per hLiMT, respectively) and when compared with the CYP3A4 activities of induced hLiMTs in wells (P>0.05). We also observed CYP3A4 activity for monoculture of 10k MSCs on-chip when no PDX cells or hLiMTs were present in culture; however, the CYP3A4 activity was lower than that of the hLiMT-MSC co-culture (**Supplementary Figure 4**).

### 2.3. On-chip and well-based prodrug-testing assay

To test ifosfamide on patient-derived samples, the PDX cells and MSCs were co-cultured with hLiMTs (**Figure 2**). The system was exposed to different concentrations of ifosfamide, as illustrated in **Supplementary Figure 2**. To compare the results achieved with our platform to those obtained with well-based assays, we tested the PDX cells in parallel with our system and in a well plate. To recapitulate liver metabolism in the well plate-based assay, we added hLiMTs to separate culturing wells with different ifosfamide concentrations and incubated them for 24 hours. The conditioned medium in hLiMT wells potentially containing metabolites was then transferred into wells with PDX cells (**Supplementary Figure 2**.a, **Supplementary Figure 5**). We decided not to co-culture hLiMTs with PDX samples in the same well to prevent direct physical interaction between hLiMTs and PDX cells or MSCs. Co-culturing of hLiMTs with MSCs in a standard well would result in a non-physiological interaction of cells due to MSCs physically attaching to the hLiMTs and partially covering their surface, which might change the CYP3A4 activity of hLiMTs and could impact the assessment of liver-mediated drug-response profiles (**Supplementary Figure 6**). Such non-physiological interactions could result in unpredictable changes in the behavior of either cell type. Cell collection for off-chip viability assessments would also be hindered by direct physical contact between hLiMTs and the MSCs or PDX cells. Moreover, we decided not to use permeable membrane inserts, e.g., transwell systems, as means to physically separate hLiMTs from the adherent MSCs or PDX cells as spheroids could attach to and spread on such inserts. The hLiMTs would then lose their 3-dimensional morphology and, potentially, their biotransformation activity ^50,51^.

We first sought to monitor the viability of PDX cells upon culturing on different materials, in different chip designs, and using different surface treatments (**Figure 4**.a). Our results showed that the donor cells of PDX-1, derived from a high-risk leukemia patient, featured a viability as high as ~80% in fibronectin-coated microfluidic culture chambers on our platform, which was similar to, although slightly lower than that in standard tissue-culture wells. To investigate the impact of microfluidic flow and on-chip culturing on PDX cells, we performed a population comparison, based on flow-cytometry scattering parameters of viable PDX-1 cells that were cultured in wells or on chips (**Supplementary Figure 7**). We did not observe any significant difference in the scattering parameters of the two populations, which indicates that our microfluidic platform did not induce morphological changes ^52,53^. We also confirmed that ifosfamide exposure did not affect MSC viability (by culturing 10k MSCs in wells at different concentrations of ifosfamide). There was no non-specific response of the tested PDX samples to the drug compound (**Figure 4**.b). We then exposed the PDX cells from the same donor to four concentrations of ifosfamide (0.01, 0.1, 1, and 10 mM) in standard wells to evaluate the cytotoxicity of the compound (**Supplementary Figure 8**). Cell viability was measured after a 3-day drug-exposure treatment by collecting the cells from each well or channel. The number of alive cells was quantified by FACS analysis after staining the dead cells with propidium iodide (PI) and by normalizing the count of viable cells to that of the control condition (no-drug treatment). Our measurements showed that a 0.01 mM ifosfamide treatment had no effect, and the PDX cells exposed to 10 mM ifosfamide showed zero viability, possibly due to the cytotoxic properties of the prodrug compound at high concentrations. As the maximum plasma concentration (C_max_) of ifosfamide at the highest clinically recommended dose has been reported to be 431 μM ^54^, we opted for excluding the 0.01 and 10 mM ifosfamide treatments from further experiments and to test prodrug efficacy at concentrations of 0.1 and 1 mM, which corresponded to clinically relevant prodrug-doses ^54^.

**Figure 4.**
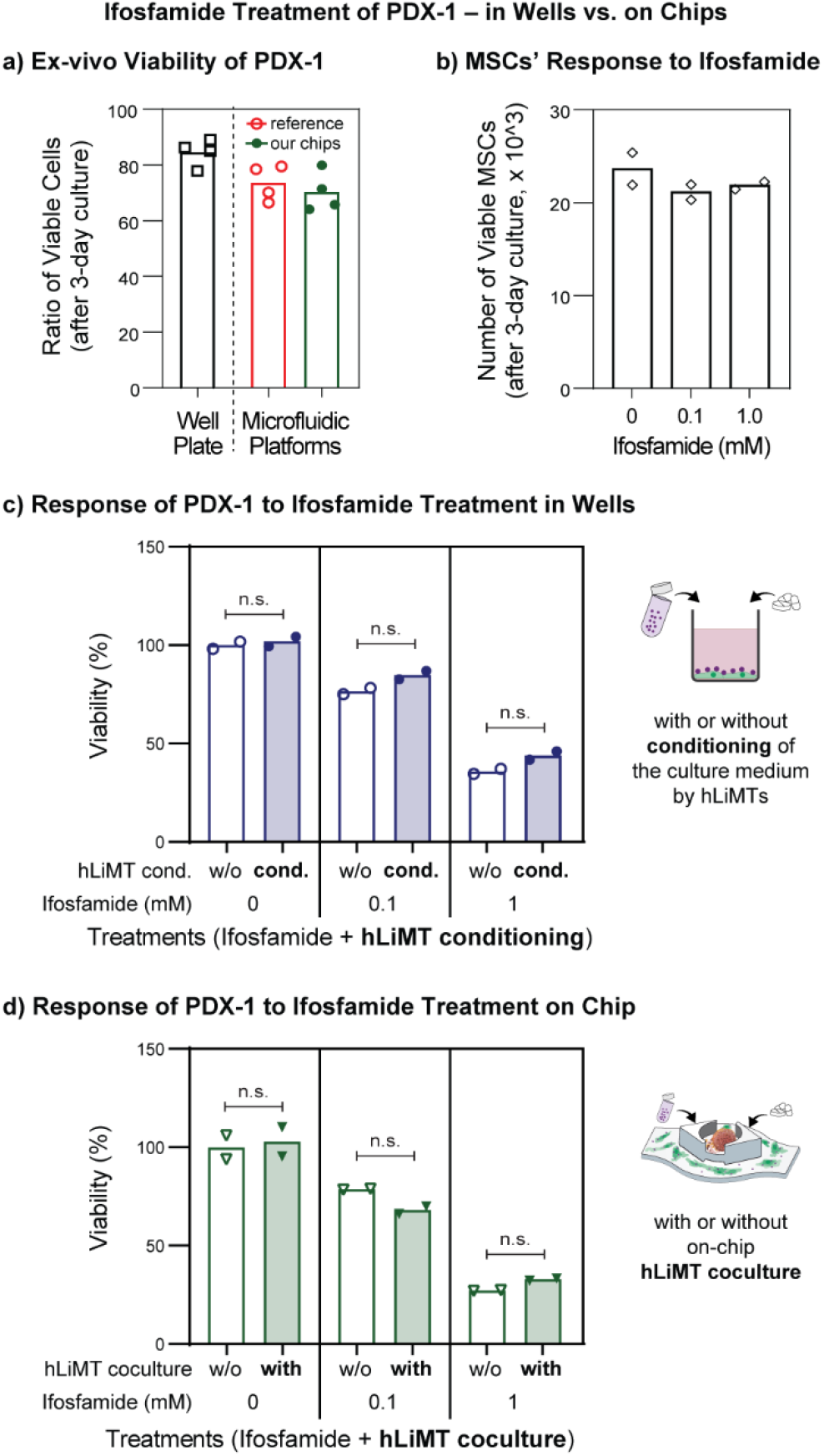
**a)** Ex-vivo viability of PDX-1 cells (ratio of viable cells) in a tissue-culture-treated 96-well plate and in microfluidic platforms (i.e., a reference polystyrene chip with no modifications, and our platform with the custom PDMS bottom and elevated hLiMT compartments). More than 70% of PDX-1 cells were viable in the microfluidic device after 3 days of culturing, similar to the standard well-based culturing. **b)** The response of MSCs’ to ifosfamide in a well plate. Ifosfamide treatment did not impact MSC proliferation in the wells. The number of cells was quantified by using fluorescent counting beads (n=2 replicates in wells). **c, d)** Normalized viabilities (as a percentage of control conditions) of PDX-1 samples after ifosfamide treatment (0, 0.1, or 1 mM) **c)** in a well plate with or without 24-h-conditioning of the culture medium and **d)** in our platform with or without on-chip hLiMT co-culturing. The ifosfamide concentrations are reported below each bar plot. After a 3-day treatment, viability decreased as a function of increasing ifosfamide concentration. The hLiMT metabolism did not cause any significant variation in PDX cell viability, neither in wells nor on chips. Cell viability in our platform was similar to that in the well plate. The plotted viabilities are calculated by normalizing the measured number of viable cells for each condition to the average number of viable cells of the controls in the wells (0 mM without liver conditioning) or the chips (0 mM without liver co-culturing). Each data point represents the mean of two technical replicates for each treatment condition in the wells or on the chips (n=2 per condition). n.s. = not significant.

We then investigated the effect of hepatic transformation of ifosfamide on cell viability of the PDX-1 cells in the well-plate assay and in our platform by implementing a co-culture with hLiMTs (**Figure 1**c). The resulting viabilities in the well plate were 101.9 ± 3.4%, 84.9 ± 2.9% and 43.9 ± 2.9% for the liver-conditioned medium in comparison to controls without liver conditioning, which featured viabilities of 100 ± 2.6%, 76.7 ± 2.1% and 35.9 ± 1.8%, for ifosfamide concentrations of 0, 0.1 mM and 1 mM (**Figure 4**.c). With our platform, we found viabilities of 102.9 ± 10.5%, 68.1 ± 2.6% and 32.9 ± 0.7% when PDX cells were co-cultured with hLiMTs in comparison to the monoculture case exhibiting viabilities of 100 ± 8.5%, 78.7 ± 0.3% and 27.2 ± 0.2%, for 0, 0.1 mM and 1 mM ifosfamide concentrations (**Figure 4**.d). Comparing the resulting viabilities of the well-based assays and our platform to their controls, neither liver-conditioned medium nor co-cultures with hLiMTs showed any detrimental effect on PDX viability for any of the tested drug concentrations (**Supplementary Table 2**). Moreover, responses to ifosfamide treatment on chips and well plates without liver-mediated metabolic effects showed consistent results between the two platforms, which confirmed the suitability of our platform for PDX-based drug testing (**Supplementary Figure 9**).

### 2.4. Identifying PDX responses to ifosfamide treatment

After confirming that our platform enabled the co-culturing of PDX leukemia cells and hLiMTs, and that hLiMTs maintained the expression of CYP enzymes for biotransformation of prodrugs, we tested the susceptibilities of four additional patient-derived samples (PDX-2 to PDX-5) to ifosfamide and metabolites in our platform (**Figure 5**). These PDX samples consisted of both, standard-risk and high-risk pediatric leukemia patients (**Supplementary Table 1**). Varied responses to ifosfamide were detected for the different PDX samples, and the addition of ifosfamide generally induced a decrease in cell viability with increasing concentration. In comparison to other samples, PDX-2 showed higher tolerance to ifosfamide treatment in our platform, and the 0.1 mM treatment did not decrease PDX viability significantly (**Figure 5**.a), whereas 1 mM treatment showed a reduction in viability by approximately 50%. PDX-2 cell viability was not altered by co-culturing with hLiMTs in the presence of the prodrug compound. For PDX-3 cells, exposure to 0.1 mM and 1 mM of prodrug solutions massively decreased the viability, independent of the presence of ifosfamide metabolites upon hLiMT transformation (29.7 ± 2.0% and 32.9 ± 8.1% for 0.1 mM exposure, 16.6 ± 4.1% and 17.4 ± 5.6% for 1 mM exposure; with and without liver co-culture, respectively) (**Figure 5**.c). PDX-4 viability was strongly affected by the exposure to ifosfamide, and very low viability (10.7 ± 2.3% with and 19.7 ± 8.0% without liver co-culture) was obtained after exposure to 1 mM ifosfamide, either with or without hLiMTs on-chip (**Figure 5**.d). Moreover, upon exposure to the 0.1 mM concentration of ifosfamide, PDX-4 viability decreased to 22.9 ± 3.7% in the case of co-culturing with hLiMTs, which was significantly lower than the 69.9 ± 23.3% viability for PDX-4 without hLiMT co-culturing (P<0.05, **Figure 5**.d). PDX-4 viabilities in well plates showed similar trends like on chip, but the distinct difference at 0.1 mM upon hLiMT co-culturing (**Figure 5**.d), as compared to culturing without hLiMTs, was not visible for conditioned/unconditioned medium in the well plate (**Figure 5**.e). Finally, PDX-5 showed low in vitro viability and the response to ifosfamide treatment could not be measured, as the low number of viable cells at the end of the assay prevented any reliable data interpretation. Therefore, PDX-5 was excluded from subsequent analysis and discussion (**Supplementary Figure 10**).

**Figure 5.**
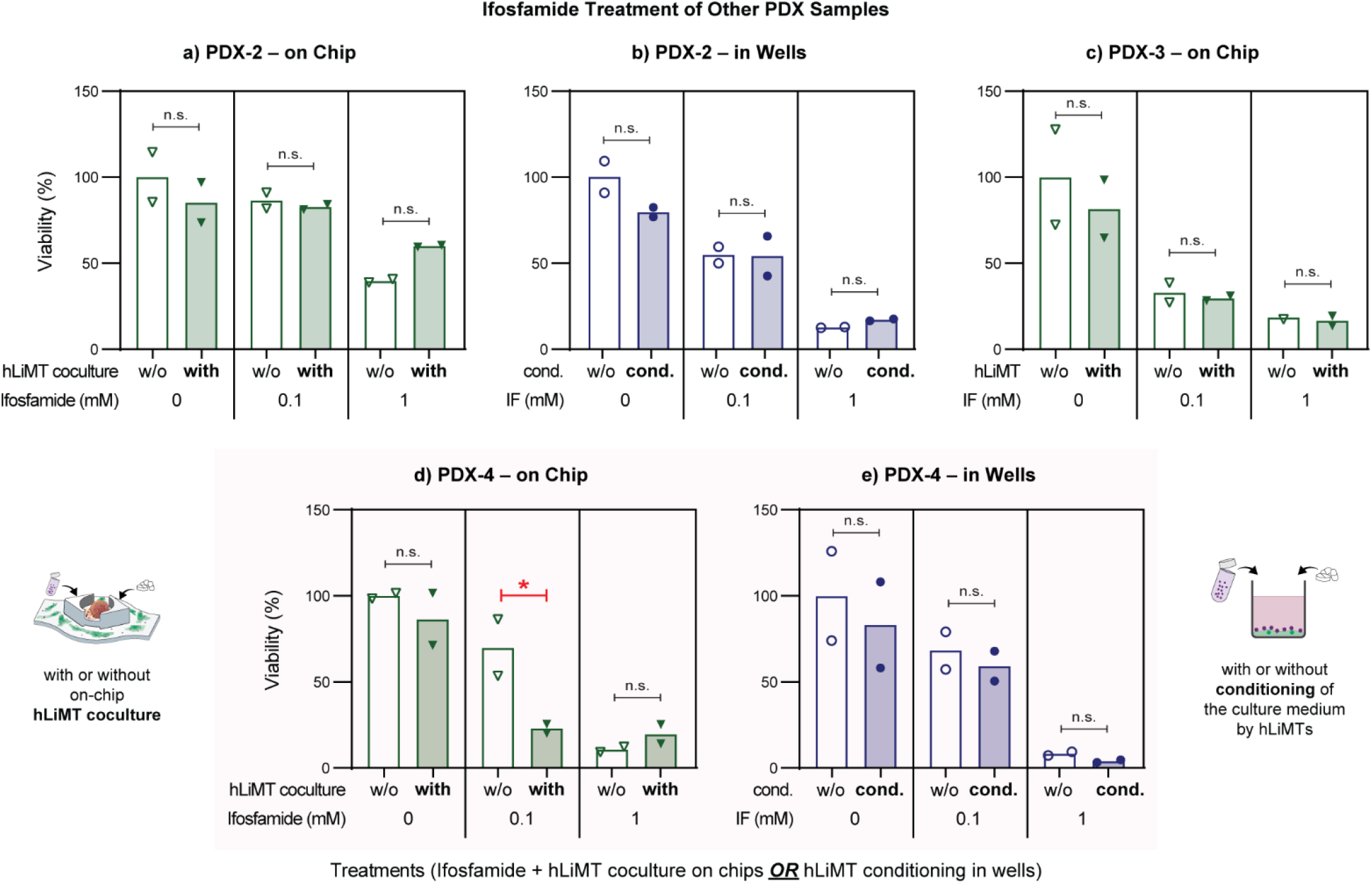
Ifosfamide treatment of PDX samples from different patients (viability given as percentage of the viability under control conditions for each PDX sample). **a)** PDX-2 did not show any effect of on-chip liver metabolic conversion of ifosfamide, and no difference in viability could be observed when the hLiMTs were present in co-culture. The tolerance of PDX-2 to ifosfamide treatment was higher in comparison to other PDX samples. **b)** The response of PDX-2 to the treatment in wells was similar to that in our platform. The sample viability decreased with increasing ifosfamide concentration, and medium conditioning did not result in a difference in the resulting response. **c)** PDX-3 did not show a response to ifosfamide metabolites when co-cultured with hLiMTs. The viability decreased with increasing ifosfamide concentration, and exposure to 0.1 mM ifosfamide resulted in a drastic decrease in viability. **d)** PDX-4 showed an effect of liver metabolism for the 0.1 mM ifosfamide treatment on chip, and the viability decreased in comparison to the ifosfamide treatment without hLiMT co-culturing. * P<0.05. **e)** Treatment of PDX-4 was repeated in a well plate. The medium preconditioning with hLiMTs did not have any effect, in contrast to the on-chip co-culturing.

To verify the role of liver metabolism in the responses observed in our platform, we compared the on-chip responses of PDX samples to the responses obtained in a well-based assay with or without medium conditioning by hLiMTs (**Figure 5**, **Supplementary Figure 11**). We used linear regression models (analysis of variance) for estimating treatment-group means and for contrast analysis to compare the treatments. A summary of the statistical analysis for PDX responses modeling all factors (hLiMT metabolism, ifosfamide concentration, and culture platform) is provided in **Supplementary Table 2**.

The responses of PDX-1 and PDX-2 to the ifosfamide treatment in well-plates were similar to their respective responses observed in our microfluidic platform: For both samples, PDX-cell viability decreased with increasing ifosfamide concentrations, and liver conditioning of the medium did not result in a significant difference in the response (**Figure 4**, **Figure 5**.a,b). However, in comparison to PDX-1, PDX-2 viability was more strongly affected by the exposure to ifosfamide, and a low cell viability (16.9 ± 0.8% with and 12.6 ± 0.4% without liver conditioning) was obtained after exposure to 1 mM ifosfamide in wells. On the other hand, PDX-3 showed low in vitro viability in the well plate, and the response to ifosfamide treatment could not be reliably measured (**Supplementary Figure 11**).

For PDX-4, exposure to 1 mM ifosfamide in the well plate was lethal to the cells, and no difference in viability was noticed upon medium conditioning. The viability of PDX-4 cells under 0.1 mM ifosfamide exposure was reduced by approximately 30% over the three-day exposure, similar to the viability decrease that was seen for the on-chip assay without hLiMTs. The well plate-based assay showed no difference in cell viability between the treatment medium and liver-conditioned medium for the 0.1 mM ifosfamide concentration. In contrast, we observed an approximately 80% decrease in cell viability for the on-chip co-culture experiment with hLiMTs as compared to PDX-4 samples without hLiMTs, which hints at a large susceptibility of PDX-4 to liver-mediated metabolites of ifosfamide.

To confirm the response of PDX-4 to on-chip hLiMT-mediated ifosfamide metabolization, we exposed the PDX-4 cells in co-culture with hLiMTs to 0.1 mM ifosfamide in presence or absence of 1 μM ritonavir that is known to inhibit CYP3A4 activity (**Figure 6**). The sample viabilities after 0.1 mM ifosfamide treatment with or without the hLiMT co-culture confirmed the previously observed PDX-4 susceptibility to ifosfamide metabolites (55.11 ± 5.2% difference between the treatment with and without the presence of hLiMTs, P<0.001). The dosage of ritonavir alone resulted in a lower sample viability in comparison to control conditions, which was similar for the presence (61.02 ± 4.6%) and absence (60.52 ± 9.8%) of hLiMTs indicating a certain toxicity of ritonavir. Nevertheless, we observed a statistically significant decrease in toxicity of ifosfamide and a higher PDX viability (15.62% ± 5.9% increase in viability) upon simultaneous dosage of ifosfamide and ritonavir in comparison to exposure to 0.1 mM ifosfamide alone in co-cultures of PDX-4 cells with the hLiMTs. The inhibition of CYP3A4 by ritonavir reduced the formation and toxic effects of ifosfamide metabolites so that PDX viability was similar to that upon ritonavir exposure alone (58.20 ± 6.5%, P>0.05).

**Figure 6.**
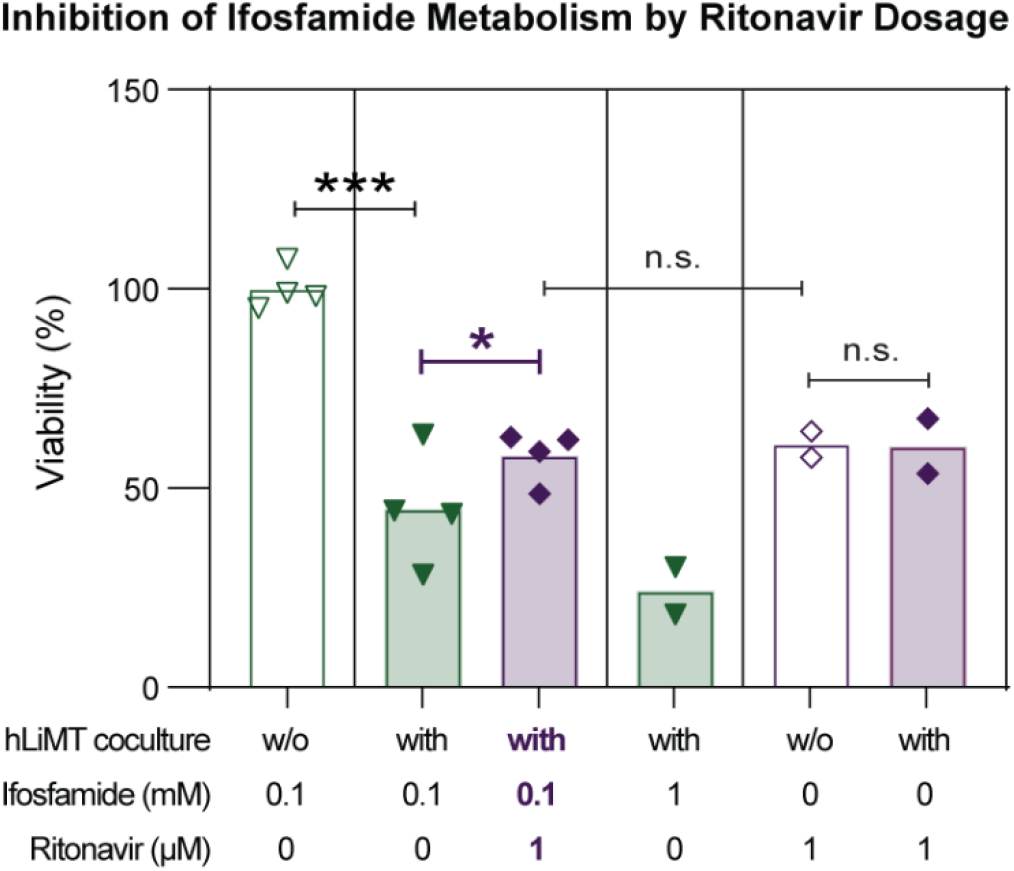
Inhibition of CYP3A4 activity by ritonavir to verify the PDX-4 response to ifosfamide metabolites on chip. Ifosfamide treatment decreased PDX-4 viability with increasing drug concentration, as in previous experiments. Comparison of 0.1 mM ifosfamide treatment with or without the hLiMTs in coculture confirmed the observed response of PDX-4 to the liver-mediated ifosfamide metabolites on chip. Simultaneous dosage of ifosfamide and ritonavir showed a statistically significant increase in PDX-4 viability when compared to treatment with 0.1 mM ifosfamide in co-culture with hLiMTs. PDX-4 viability upon the simultaneous dosage of ifosfamide and ritonavir showed no significant difference to the condition when exposed to ritonavir alone, either with or without hLiMT coculture. n=2 or 4, * P<0.05, *** P<0.001, n.s. = not significant).

## 3. Discussion

In an effort to identify therapeutic strategies for improving the survival rate for leukemia patients, drug screening using patient-derived leukemia cells has enabled measuring the susceptibility of primary leukemia cells to common anticancer compounds and repurposed drugs. However, in vitro drug screening has usually been limited to the detection of direct compound effects. Complex drug effects, such as liver-mediated metabolic conversion of compounds such as prodrugs are only rarely assessed. Therefore, we developed a microfluidic drug-testing platform for co-culturing of primary leukemia cells and human liver microtissues (hLiMTs), which was aimed at including hepatic transformation of prodrugs into the overall efficacy testing.

We used hLiMTs, formed from primary hepatocytes of multiple donors, to recapitulate the CYP-mediated metabolism of prodrugs in vitro (**Figure 3**). The use of multi-donor hLiMTs provides consistency over multiple experiments in comparison to single-donor tissues, which usually feature high inter-donor variability due to varying donor-cell CYP activities ^55–57^. Only consistent liver-metabolism parameters across the different experiments enable to identify the sensitivity of PDX cells to prodrug treatments, and to compare the observed response profiles with each other. For treatment, however, patient-specific responses inevitably depend on the patients’ individual metabolic activities ^57^. Therefore, the use of patient-derived liver microtissues could potentially increase the predictive power of the assay by recapitulating the patient-specific response in vitro. Such an approach would require additional sample material from the patient, namely mature hepatocytes or hepatocytes derived from pluripotent stem cells, that then could be used for the formation of functional microtissues.

The metabolic activity of liver microtissues in vitro has been extensively reported in literature, both for static well-based conditions and dynamic perfusion culture conditions ^39,40,56^. Our mass-spectrometry measurements showed that the hLiMTs were able to metabolize ifosfamide and that the biotransformation of the prodrug was driven by the liver-specific CYP enzymes. The capacity of hepatic biotransformation could further be increased by 24-h induction of the hLiMTs in 1mM ifosfamide solutions prior to exposing the PDX cells to the drug compound. This procedure allowed us to avoid high prodrug concentrations (higher than 1 mM), which were shown to affect PDX viability. In an earlier study, we had quantified the metabolic transformation of the prodrug midazolam by hLiMTs. Our results evidenced that the CYP3A4 activity of the hLiMTs remained stable over a culture period of 14 days and that the activity was comparable for microtissues under static well-plate conditions and under perfusion conditions ^33^ similar to the conditions in this study. Moreover, we assessed the CYP activity of the hLiMTs for static, well plate-based cultures and for on-chip co-culturing conditions by using a biochemical assay based on the metabolic transformation of Luciferin-IPA to D-Luciferin. Luciferin-IPA is metabolized mainly by CYP3A4 ^58^, following phase I metabolic pathways that are similar to that of ifosfamide ^24^. The obtained results confirmed the ability of the hLiMTs to metabolize ifosfamide in our platform. CYP enzymes, especially CYP3A4, have been reported to be expressed by bone marrow stroma and are thought to play a key role in clinical drug resistance through chemoprotection ^59,60^. In line with this concept, we observed CYP3A4 activity by the MSC feeder layer when cultured alone on-chip, however, 2-fold lower than the CYP3A4 activity of hLiMTs (**Supplementary Figure 4**).

In this proof-of-principle study, we tested our platform with 5 PDX samples from standard- and high-risk pediatric ALL patients (**Supplementary Table 1**). We characterized the compatibility of the microfluidic platform and established the on-chip prodrug testing assay with ifosfamide using the PDX-1 sample. The additional four PDX samples (PDX-2, PDX-3, PDX-4, and PDX-5) were then used to test ifosfamide treatment on chip with the established assay. One sample, PDX-5, showed low in-vitro viability and was, therefore, excluded from further analysis. Ifosfamide treatments of all other PDX samples were carried out in parallel in wells to elucidate the role of short-lived metabolites in the response profiles of these samples. The PDX-3 sample showed low viability in the well-plate assay and was excluded from further analysis.

A comparison of PDX-1 responses without liver-mediated transformation of ifosfamide, i.e., without liver medium conditioning or on-chip hLiMT co-culturing, between the standard well-plate assay and on-chip testing confirmed that the culturing of PDX cells in our microfluidic platform did not affect or alter cell viability or the outcome of the drug-response-profiling experiments (**Supplementary Figure 9**, **Supplementary Table 2**). Cell viability was measured after three days in culture, both on-chip and in standard well plates, using FACS analysis. No difference in relative PDX viability between PDX/MSC and hLiMT co-cultures (or medium conditioning in case of the well-plate assay) and PDX/MSC monocultures could be observed under control conditions, which showed that hLiMTs did not affect PDX viability. The obtained results indicated that PDX-1 did not show a pronounced response to ifosfamide metabolites. Although the absolute number of viable cells in the microfluidic platform after 3 days was lower in comparison to that in the well plate, the relative viabilities and measured responses of PDX-1 cells showed very good agreement between the two culturing methods for all tested ifosfamide concentrations **(Figure 4)**. The agreement in experimental outcome between the two assay types, microfluidic platform and well plate, proved that our assay could replicate drug responses obtained with traditional well-based assays.

Upon testing the other PDX samples on our platform (**Figure 5**, **Supplementary Figure 10**), we observed varying viability. Such variation between PDX samples has already been reported in the literature as optimal culture conditions are patient-dependent and may, therefore, differ for PDX samples from different patients ^8,11^. Two of the samples, PDX-2 and PDX-3, did also not show a detectable response to liver-induced ifosfamide metabolites upon treatment in co-culture with hLiMTs, similar to PDX-1. However, ifosfamide exposure decreased the viability of PDX-1, PDX-2, and PDX-3 to a different extent upon increasing concentration, which suggests that PDX samples of individual patients feature different susceptibilities to the drug compound. In contrast, the PDX-4 sample in co-culture with hLiMTs showed a much lower viability value upon exposure to 0.1 mM ifosfamide in comparison to exposure in a monoculture without hLiMTs. The drastic decrease in cell viability suggests a high susceptibility of PDX-4 to ifosfamide metabolites, produced by the hLiMTs on-chip, which is clearly different from the results obtained for the other PDX samples (**Supplementary Figure 12** shows a direct comparison of on-chip dose-response curves of two samples showing different prodrug susceptibility, namely PDX-1 and PDX-4). In the wellplate assay, there was no difference in PDX-4 cell viability for exposures to 0.1 mM ifosfamide with and without medium-conditioning by hLiMTs, which suggests that the presence and toxic effects of ifosfamide metabolites could be probed in our platform but not in the well plate. Furthermore, simultaneous dosage of ritonavir and ifosfamide resulted in a decreased toxicity of ifosfamide for PDX-4 cells, as ritonavir inhibited the CYP3A4 activity of hLiMTs. The markedly lower viability of PDX-4 cells upon exposure to 0.1 mM ifosfamide in co-culture with hLiMTs, and the increase in viability upon co-dosing ritonavir provides evidence that the PDX-4 cells specifically react to highly toxic metabolites of ifosfamide, produced by the liver microtissues, such as isophospharamide mustard ^24,48^. These metabolites are known to be unstable and short-lived in aqueous solutions and decay within a few hours ^24,27^. Only the direct interaction and co-culturing with hLiMTs in the same liquid phase in our platform yielded an increased cytotoxic effect, as the short-lived, liver-metabolism-induced prodrug metabolites were continuously produced, while the conditioning of medium through hLiMTs and a subsequent transfer to target PDX cells did not yield this cytotoxic effect. At high drug concentrations, MSCs may contribute to reducing PDX viability by transforming ifosfamide through self-expressed CYP enzymes. However, such high concentrations would exceed the reported maximum plasma concentration (C_max_) of ifosfamide in patients at clinically recommended doses ^54^ and are, therefore, of limited clinical interest.

During static culturing, cells may suffer from nutrient or drug depletion over time, as there is no fluid flow and the transport of molecules is limited by diffusion ^31^. To avoid this issue, we operated our platform on a tilting device to generate a pump- and tubing-free flow by inducing a pressure difference between the two reservoirs flanking the microfluidic channels ^22,32^. The resulting gravity-driven flow ensured continuous recirculation and redistribution of metabolites produced by the hLiMTs and of nutrients in the culture medium and provided continuous interaction between the diverse cell types through signaling molecules in the liquid phase.

In the microfluidic chips, PDX samples typically featured lower viability in comparison to the well-based assays (**Figure 4**.a). The comparison of flow cytometry data of PDX cells (e.g., side scattering and forward scattering parameters related to granularity or intracellular complexity and size of cells, and CTV-fluorescence amplitude), cultured on the two platforms, did not exhibit any significant differences, which implies that there was no perceivable impact of culturing and flow conditions on the morphology of PDX cells (**Supplementary Figure 7**). On the other hand, the well-plate-based and on-chip responses of several PDX samples for which the liver metabolism did not play a role were very similar, which evidences that our platform did not generally affect or alter the response profiles of PDX cells (**Supplementary Figure 9**). We suspect that somewhat lower on-chip viabilities could be a consequence of tedious cell retrieval from the less accessible channel regions of the microfluidic chips. Potentially stressful washing and cell retrieval steps were required to collect the cells from the microfluidic channels, which was not the case for standard well plate assays. Moreover, the experimental protocol required a multiplicity of manual pipetting steps to prepare, seed, collect and analyze samples with relatively long waiting times in between, which limited the overall throughput of the study.

All the PDX cells used in this study were provided in anonymized cryopreserved vials of 10-50 million PDX cells derived from ALL patients. The samples’ risk stratification was based on the minimal residual disease (MDR) analysis of the patients that the PDX cells originated from. As ALL patients are usually treated with a cocktail of compounds to maximize treatment efficacy, it is very difficult to directly compare the results obtained here with patient’s treatment responses. Nevertheless, the screening of patient primary leukemia cells with potential chemotherapeutic agents after diagnosis has been reported to improve clinical outcomes ^11^.

Our platform enables the inclusion of prodrugs and the corresponding short-lived metabolites in functional precision-medicine assays. In addition, co-cultures with metabolically active liver models during drug testing opens a route to investigating other metabolism-induced effects during drug administration. For instance, a system similar to our microfluidic platform was used to recapitulate in vitro drug-drug interaction phenomena and how those can cause drastic variations in drug efficacy ^33^. Cancer patients are usually treated with a combination of anti-cancer compounds ^2,3^. Our platform could be used for testing of such combination therapies on patient-derived or primary samples from high-risk ALL patients. Advanced microfluidic systems could be used for establishing personalized treatments by including hepatic metabolism and the co-administration of multiple drug compounds into drug testing and screening. Such systems, including parts of our platform, are often realized in polydimethylsiloxane (PDMS), a soft polymer material that features high biocompatibility, optical transparency, and allows for rapid prototyping ^62,63^. However, PDMS is also known to absorb small hydrophobic molecules, which limit its use for drug testing application ^64,65^. Therefore, before translation to clinical settings, these systems need to be realized in inert materials, such as standard hard plastic materials, commonly used for drug testing and amenable to high-throughput fabrication.

## 4. Conclusions

We developed and tested a physiologically relevant platform that enables testing of compounds including prodrugs that produce short-lived or unstable metabolites with patient-derived leukemia cells. Our platform allows for ex-vivo testing of patient-derived leukemia samples in co-culture with and in close vicinity to human liver microtissues in the same microfluidic network. As it offers the possibility to test liver-mediated activation of prodrugs, the developed platform offers great potential for translation to clinical settings for the selection and optimization of treatments for leukemia patients. This holds particularly true for the broad use of prodrugs, such as oxazaphosphorines that are characterized by short-lived metabolites, in leukemia treatments.

While the current chip design allows for testing of up to eight conditions per plate in parallel, the throughput can be increased by modifying the microfluidic network so as to reduce the culture and sample volume. Such design modifications would enable testing a larger number of conditions without requiring larger PDX cell numbers. Additional optimization of measurement protocols and modifications of the microfluidic chip design could enable automated imaging-based cell viability assessment by high-resolution microscopy to shorten the analysis time and increase throughput.

## 5. Materials and Methods

### 5.1. Cell Culture

PDXs were generated by injecting in NSG mice primary human ALL cells, recovered from bone marrow aspirates of patients in the ALL-BFM 2000, ALL-BFM 2009, and ALL-REZ-BFM 2002 studies (informed consent was given according to the Declaration of Helsinki and ethics commission of the Canton of Zurich, approval number 2014-0383), as previously described ^14^. The samples were classified as standard-risk (SR) or high-risk (HR) according to the ALL-BFM 2000 stratification (**Supplementary Table 1**) ^66^. PDX cells were cryopreserved as anonymized vials of 10 to 50 million PDX-ALL cells. The PDX cells were thawed and seeded into the microfluidic channels or in the wells to start the drug exposure tests. More information on the thawing, seeding, and medium conditions for the PDX ALL cells is reported below.

Primary human liver microtissues (hLiMTs, 3D InSight™ Human Liver Microtissues of multi-donor primary human hepatocytes, XL size production MT-02-302-11, InSphero AG, Schlieren, Switzerland) were cultured using 3D InSight™ Human Liver Maintenance Media - AF (InSphero AG) in Akura96 Ultra-Low Attachment plates (InSphero AG). For PDX-testing experiments, the hLiMTs were used within 3 days of delivery. The hLiMTs used in this study had an average diameter of around 325 μm ± 5% (the average diameter after tissue reformation, based on the certificates of quality for hLiMT plates of each experiment).

GFP-expressing, hTERT-immortalized primary human bone marrow mesenchymal stromal cells (MSCs) ^12,18,19^ were kindly provided by D. Campana (St Jude Children’s Research Hospital, Memphis, Tennessee). MSCs were maintained in RPMI 1640 medium (Sigma-Aldrich, Buchs, Switzerland) containing 10% fetal bovine serum (Sigma-Aldrich), 2 mM L-glutamine (Thermo Fisher Scientific, Reinach, Switzerland), 100 IU/mL penicillin/streptomycin, and 1 μM hydrocortisone (Sigma-Aldrich).

### 5.2. Microfluidic Platform Fabrication and Operation

#### 5.2.1. Platform Assembly

The drug-screening platform was developed on the base of an injection-molded, microfluidic multi-tissue culture chip made of polystyrene, which had been previously designed by our group in collaboration with InSphero AG (AkuraFlow chips) ^22^. The microfluidic channels of the chip were modified so as to realize elevated compartments for culturing the hLiMTs and reduce the direct physical interaction of PDX cells and MSCs with the hLiMTs. The elevated platforms were realized by bonding a modified PDMS bottom to the polystyrene chip as illustrated in **Supplementary Figure 1**. We designed the master mold for the PDMS bottom parts on AutoCAD (Autodesk Inc., Mill Valley, CA, USA); transparency photomasks were printed by Micro Lithographic Services Ltd (Chelmsford, UK). The master mold was fabricated on a 4-inch silicon wafer with SU8-3025 by using standard photolithography steps. The thickness of the SU8-3025 structures was 20 μm. The master mold was then silanized in a vacuum desiccator with Trichloro(1H,1H,2H,2H-perfluorooctyl)silane (Sigma-Aldrich,) for 2 hours. 1:10 PDMS (Sylgard® 184, Sigma-Aldrich) was cast on the master mold and cured at 80°C for 2 hours. Cut-outs of the cured PDMS slab, which then formed the bottom of our platform, were cleaned from debris using scotch tape (4.977.763, Lyreco, Dietikon, Switzerland) and stored until assembly with the polystyrene chips.

To prepare the polystyrene parts for assembly, they were cleaned with isopropyl alcohol (IPA) and ethanol and then dried with pressurized air. Next, the chips were treated with oxygen plasma for 30 seconds at 50 W power (Harrick Plasma PDC-002, Harrick Plasma, Ithaca, NY, USA) and subsequently submerged in a solution of 2% bis[3-(trimethoxysilyl)propyl]amine (Sigma-Aldrich), and 1% DI-water in IPA at 80°C for 20 minutes following previously published protocols ^67^. The chips were then rinsed with IPA, dried in a convection oven at 70°C for 30 minutes, and then immersed in 70% ethanol in water until bonding to the PDMS bottom parts.

The PDMS bottom parts of the system with the tissue-compartment elevations were treated with oxygen plasma for 30 seconds at 50 W power, aligned, and bonded to the polystyrene chips that were silanized in the 2% bis-amino silane solution by using a custom chip aligner made of micromanipulators and a stereomicroscope. The assembled chips were then dried and cured at 70°C for 2 hours.

#### 5.2.2. Platform Operation on Tilting Device

Following the assembly, the chips were sterilized by spraying with IPA and by exposing them to UV-light for 8 hours in a sterile laminar flow hood. For the experiments, we inserted three of our platforms in rectangular holders with ANSI standard well plate dimensions that could accommodate 4 devices. To avoid medium evaporation during the experiments approximately 4 ml PBS were filled into the well in the fourth position and the chips were sealed with adhesive films featuring holes to facilitate air exchange.

On-chip flow during drug exposure assays was generated by periodically tilting the chips by a ± 3° angle, with a transition time of 30 seconds from each tilting-end position to the rest position (0° angle) and vice versa, and a waiting time of 10 minutes at each tilting-end position. Four 4-well rectangular dishes could be operated with a single tilting device (GravityFlow, InSphero AG, see **Supplementary Figure 3**).

### Experimental Procedure for Drug Response Profiling

#### 5.2.3. Culture and Treatment Media

Serum-free AIM-V medium (12055091, Gibco™ AIM V™ Medium, Thermo Fisher Scientific) was used for culturing the samples and for preparing the drug solutions. Ifosfamide (53358-100MG, Ifosfamide analytical standard, Sigma-Aldrich) stock solutions at 10 mM in AIM-V were freshly prepared and diluted to the dosing concentrations (0.1, 1, and 10 mM) for each experiment. The different ifosfamide solutions were used for hLiMT induction, for generation of the liver-conditioned medium, and as treatment solutions. All medium and coating solutions were warmed up and kept in a warm water bath at 37°C for at least 12 hours before starting the assays to prevent air bubble formation in the microfluidic channels.

#### 5.2.4. Chip and Well Plate Preparation

To promote the adhesion and growth of MSCs on our platform, the channels were coated with 100 μg/mL fibronectin (FC010, Sigma-Aldrich) in DPBS (Thermo Fisher Scientific). 100 μL of the fibronectin solution were added to the chip and incubated for 90 minutes at room temperature. After incubation, the channels were flushed with 100 μL PBS and rinsed three times with DI water. The chips were then stored overnight at 4°C. The morning after, PBS was exchanged with 100 μL of AIM-V medium at 37°C, and the chips were kept in a cell-culture incubator at 37°C, 95% humidity and 5% CO_2_ for at least 1 hour until the cell seeding.

#### 5.2.5. Seeding MSCs (Day -2)

Each channel or well was seeded with 10k MSC cells. To promote a uniform distribution of cells in the microchannel, MSCs were suspended at 1 million cells/mL in AIM-V media, and 5 μL of suspension were injected into the second and ninth microtissue compartments. On the well plates, 100 μL of AIM-V medium and 10 μL of the prepared cell suspension was added to each well (Thermo Scientific Nunc™ F96 MicroWell™ Polystyrene Plate, Thermo Fisher Scientific). The seeded chips and well plates were incubated at 37°C, 95% humidity and 5% CO_2_ for 24 hours.

#### 5.2.6. Labeling PDX Cells with CellTrace Violet and Seeding (Day -1)

PDX cells were thawed by placing the vials in a cell-culture incubator at 37°C for 5 minutes and by re-suspending the cells in 5 mL medium. To simplify the FACS analysis at the end of the experiment, PDX cells were labeled with CellTrace Violet dye (CTV; C34557, Thermo Fisher Scientific) according to the manufacturer’s protocol. The cells were re-suspended in 12.5 μM CTV solution in PBS and incubated for 20 minutes at 37°C and protected from light. The suspension was then centrifuged for 5 minutes at 200×g, and PDX cells were suspended to a concentration of 10 million cells/mL in AIM-V.

To seed cells in the microfluidic channel, a total of 10 μL of the prepared suspension (100k cells) was added through the inlets of the second and ninth microtissue compartment, as for the MSC seeding procedure. To seed cells in the well plate, 10 μL of the prepared suspension were added to the wells. The initially seeded number of PDX cells for each sample in this study is reported in **Supplementary Table 1**. The seeded chips and well plate were then placed in a cell-culture incubator at 37°C, 95% humidity and 5% CO_2_ for 24 hours before starting the drug treatment and tilting.

#### 5.2.7. Liver-conditioned medium and hLiMT induction (Day -1)

For well plate-based experiments, we conditioned the medium by adding hLiMTs to solutions of 0, 0.1, or 1 mM ifosfamide in AIM-V before treatment of the PDX samples. To be able to compare the well-based screening with the on-chip testing, we used identical medium volumes and numbers of hLiMTs for the two conditions, namely 4 hLiMTs per 200 μL solution per well condition (see **Supplementary Figure 2** and **Supplementary Figure 5**). Two hLiMTs were cultured in each well on an Akura96 well plate for 24 hours in 100 μL ifosfamide solution at different concentrations under standard mammalian cell-culture conditions.

To increase the expression of CYP enzymes, the hLiMTs to be used in co-culture with PDX samples on the chips were induced by culturing them in 1 mM ifosfamide solution in AIM-V for 24 hours under standard mammalian cell-culture conditions.

#### 5.2.8. hLiMT Loading and Starting of Drug Assay (Day 0)

On the day of drug treatment, four hLiMTs were transferred to the four center microtissue compartments of each designated device co-culture channel. The reservoirs and microtissue compartments were then sealed by using custom-cut, pressure-sensitive foils (Z734438, ThermalSeal RTS™ Sealing Films, Sigma-Aldrich) with holes for air exchange and to enable access to the reservoirs for medium exchange, sampling, and drug dosing.

For well plate-based experiments, we exchanged the medium in the target wells that hosted the PDX cells and MSCs with 200 μL of conditioned treatment medium, i.e., 100 μL from two separate liver wells (**Supplementary Figure 2**, **Supplementary Figure 5**). For the standard PDX exposure assay, we transferred 20 μL of 10x stock solutions of ifosfamide in AIM-V into the PDX wells on the treatment well plate and added fresh AIM-V medium to reach a total volume of 200 μL in the well.

For the on-chip assay, we added a total of 20 μL of 10x ifosfamide solutions to the medium reservoirs and added fresh AIM-V medium to reach a total on-chip medium volume of 200 μL. The chips were then loaded onto the GravityFlow tilting device and operated for 72 hours at standard mammalian cell-culture conditions.

### 5.3. Flow Cytometry Analysis (Day 3)

After 72 hours of treatment, the tilting was stopped, and the chips and wells were removed from the incubator. After collection of the supernatant, each channel and well was washed with 50 μL PBS. Afterward, 100 μL AccuMAX cell detachment solution (Sigma-Aldrich) was added and the chip was incubated at room temperature for ~20 minutes. Using AccuMAX for retrieving cells from the microfluidic channels and from the wells ensured retrieval of all cells and shortened the analysis time by preventing cell aggregation in the flow cytometer without decreasing viability. The cell suspensions were collected, and channels and wells were washed with 50 μL PBS. All washing fractions and the cell suspensions of each well were pooled to ensure complete retrieval of cells.

In a V-bottom well plate, 25 μL of fluorescent counting beads (1 million/mL, 424902, Precision Count Beads, BioLegend, San Diego, California USA) were mixed with 100 μL of cell suspension for each condition, and 1 μL propidium iodide solution (P4864, Sigma-Aldrich, Buchs, Switzerland) was added. For each sample, we performed two technical replicates.

FACS analysis was conducted by using a BD LSRFortessa Cell Analyzer with a high throughput sampler (HTS, BD Biosciences, Allschwil, Switzerland). The recording thresholds and voltages for forward- and side-scattering channels were adapted to the scattering amplitudes of the PDX cells individually for each PDX sample and not to the scattering amplitudes of the MSCs. Five to ten thousand events were recorded per well. CTV (for gating PDX cells) was excited at 405 nm and detected with a 450/50 band-pass filter for emission. GFP (for filtering MSCs and counting fluorescent beads) was excited at 488 nm and detected with a 505 nm long-pass filter, and a 530/30 band-pass filter for emission. PI (for quantifying dead PDX cells) was excited at 561 nm and detected with a 600 nm long-pass filter, and a 610/20 band-pass filter for emission. Flow cytometry data were analyzed by using the FlowJo software (FlowJo, LLC, Ashland, Oregon, USA). The analyses were normalized with respect to the number of counted fluorescent beads and corrected for the total volume in the wells or on chips. The total number of viable cells after 3-day incubation in wells and on chips are reported for each PDX sample in **Supplementary Table 1**. A workflow for flow cytometry analysis of PDX cells is provided in the **Supplementary Figure 13**.

### 5.4. Mass Spectrometry for Ifosfamide Metabolism

To investigate hepatic bioactivation of ifosfamide, hLiMTs in Akura96 well plates were first induced by exposure to 1 mM ifosfamide in AIM-V. Then, the wells were washed and 25 μL of 1 mM ifosfamide solution in AIM-V were added with or without ritonavir (1 μM, Sigma-Aldrich). 5μL supernatant solutions were collected after 0, 24, 48, and 72 h from individual wells and snap-frozen into AIM-V (45 μL) on dry ice. All samples were frozen and shipped to Admescope Ltd. (Oulu, Finland), where they were semi-quantitatively analyzed by ultra-performance liquid chromatography-mass spectrometry (UPLC-MS). For the mass spectrometry measurements presented in this study, the estimated limit of detection of alco-ifosfamide was 100 nM and that of 2-dichloroethylifosfamide was 200 nM. The plotted data were normalized with respect to the signals of a 1 mM ifosfamide solution measured at the same time intervals with mass spectrometry to compensate for any evaporation effects.

### 5.5. CYP3A4 Assay

CYP3A4 activity of hLiMTs in monoculture or in co-culture with MSCs was assessed using the P450-Glo CYP3A4 Assay (Promega, Dübendorf, Switzerland) with luciferin isopropyl acetal (luciferin-IPA) as substrate. The activity of individual hLiMTs was measured in static well plates, while the activity of 4 hLiMTs in monoculture or in co-culture with MSCs was measured for in the microfluidic platform and under perfusion.

### 5.6. Microscopy Imaging

Bright-field images of MTs in well plates were acquired using a Cell3iMager Neo cc-3000 (Screen Holdings Co., Ltd, Kyoto, Japan) to track the loading and retrieval of hLiMTs. Bright-field and fluorescence images of cells and MTs in the microfluidic platform were acquired using an inverted microscope equipped with a CrestOptics X-Light v3 spinning disk confocal (Nikon Eclipse Ti2-E; Nikon Europe B.V., Amsterdam, Netherlands). The images were processed by using ImageJ.

### 5.7. Data Analysis and Statistics

Experimental results are generally presented by bar plots for treatment group means together with corresponding biological replicates (*n=2* or as stated in the figure captions), each averaged over 2 technical replicates (same sample recorded twice by the flow cytometry analyzer). CYP3A4 activities were compared with each other by performing an unpaired t-test with Welch’s correction using GraphPad Prism.

To analyze the FACS data of the PDX samples, we used an ANOVA (analysis of variance) model of the form *log(viability)~ifosfamide*hLiMT*platform*. Here, *viability* is given as percentage, i.e., the ratio of the number of viable cells after each treatment in relation to that of the control treatment; *ifosfamide* is the concentration of ifosfamide at levels ‘0’, ‘0.1’ and ‘1’; *hLiMT* encodes the liver metabolism (hLiMT co-culturing or conditioning) at levels ‘with’ and ‘without’; and *platform* encodes the culture platform used with levels ‘chip’ and ‘well’. Using this model allowed us to pool errors over experimental conditions and provided 12 degrees of freedom for treatment comparisons (compared to only 2 degrees of freedom when directly comparing naïve-specific conditions). Based on the fitted model, we estimated marginal means and calculated linear contrasts between treatment group means using the *emmeans(*) package in R (version 1.8.0). For PDX-3, for which in-well results were not included in the discussion, we used a reduced model, *log(viability)~ifosfamide*hLiMT* (df=6). In our analyses, we performed multivariate adjustments and assumed the same residuals across the family of conducted pairwise tests. The statistical analyses of PDX samples and some key contrasts between treatment factors are summarized in **Supplementary Table 2**.

### 5.8. Data Availability

The experimental data presented and discussed in this work are available within the paper and in the Supplementary Material. The analysis data generated for this study are available on reasonable request.

## Author Contributions

**F.G.** designed and fabricated the microfluidic devices, designed, and performed all experiments, analyzed and interpreted data, and wrote the manuscript. **A.K.** conceived the project, ran initial validation experiments, and edited the manuscript. **C.L.** provided experimental advice, contributed to the MS experiment and flow cytometry analysis, and edited the manuscript. **M.G.** contributed to the initial validation experiments with A.K. **H.-M.K.** contributed to experimental design and statistical analysis of the results. **K.R.** conceived the project, initiated experimental realization, and supervised initial experiments. **B.B.** conceived the project, provided cell culture, patient-derived samples, and clinical-relevant advice, and edited the manuscript. **A.H.** provided experimental advice and led the experimental project, edited the manuscript. **M.M.** supervised the project and provided experimental advice, contributed to platform and experimental design, and the MS experiment, and edited the manuscript. All authors discussed the results and implications and commented on the manuscript at all stages.

## Acknowledgments

This work was financially supported by the Personalized Health and Related Technologies (PHRT) initiative of the ETH Domain (Project No. SFA-PHRT-2017-309). We acknowledge Dr. Olivier Frey, and InSphero AG for providing the Akura™ Flow polystyrene chips and human liver microtissues throughout the experiments, and Jana Petr for their support with confocal imaging of the cell cultures. Further, we thank the flow cytometry team at the D-BSSE Single Cell Facility for their on-site assistance and guidance with analyses. Finally, we acknowledge current and previous members of the Bio Engineering Laboratory at ETH Zurich, and of the Pediatric Oncology Research Group at University Children’s Hospital Zürich for their valuable discussions and input on the conducted work.

