## Supplementary Material for "Microphysiological Drug-Testing Platform for Identifying Responses to Prodrug Treatment in Primary Leukemia"

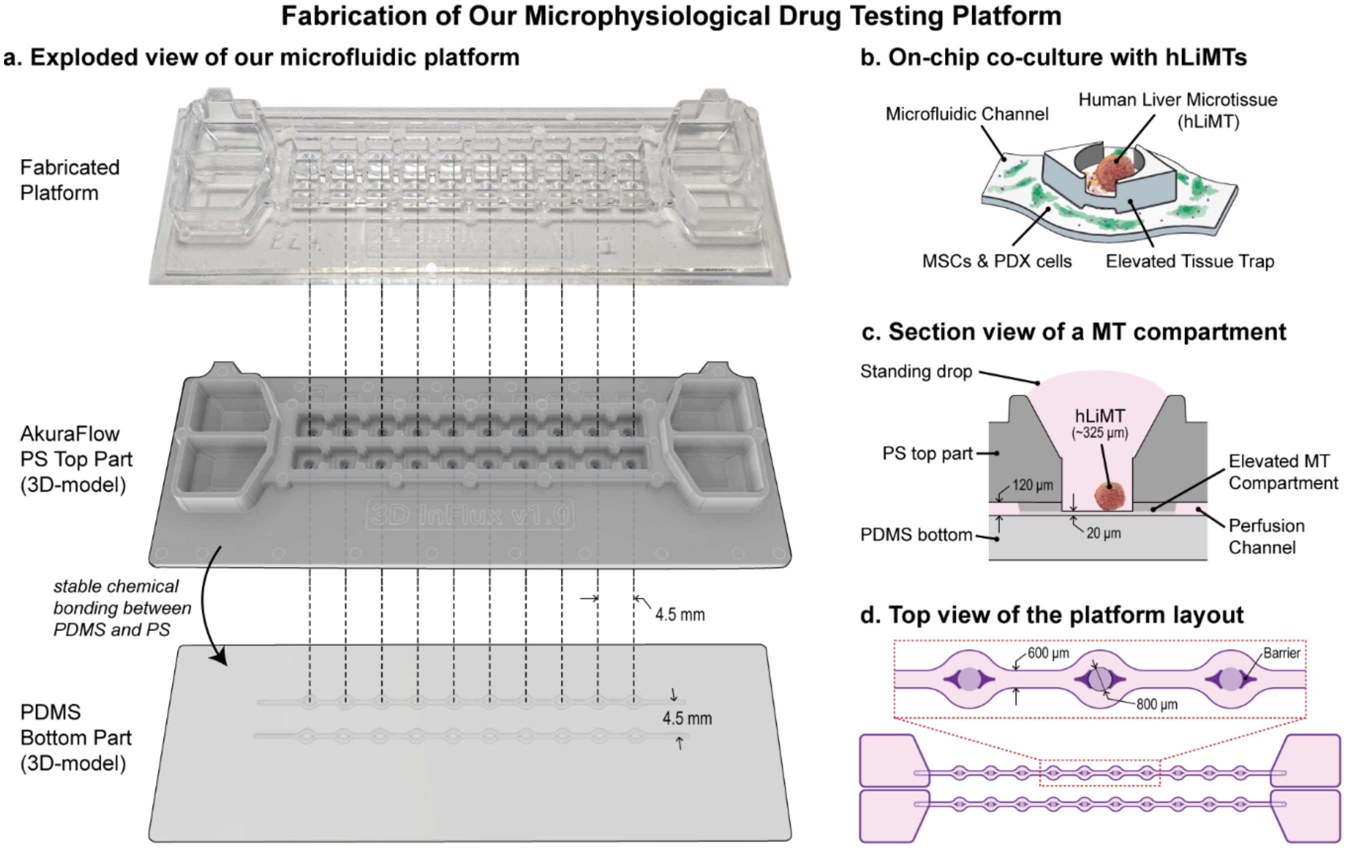


***Supplementary Figure 1*** *Fabrication of the microphysiological drug testing platform, showing the standard SLAS 384-microwell-plate spacings between the two channels and between the microtissue compartments.* ***a)*** *Photograph of the fabricated platform, and exploded view of the platform showing the polystyrene top part (commercially available as AkuraFlow chip) and the PDMS-based bottom part with the additional 20-µm-high microfluidic channel and plateau structures facing the microtissue inlets in the polystyrene chips to realize elevated microtissue compartments.* ***b)*** *Illustration showing the diverse cell types (MSCs in green, PDX cells in violet and a hLiMT) cultured in and around the elevated microtissue compartment.* ***c)*** *Cross-sectional view of one microtissue compartment and of the interconnecting microfluidic channel showing the channel and microtissue dimensions.* ***d)*** *Top and close-up views of the microfluidic channel layout showing channel and microtissue compartment dimensions.*

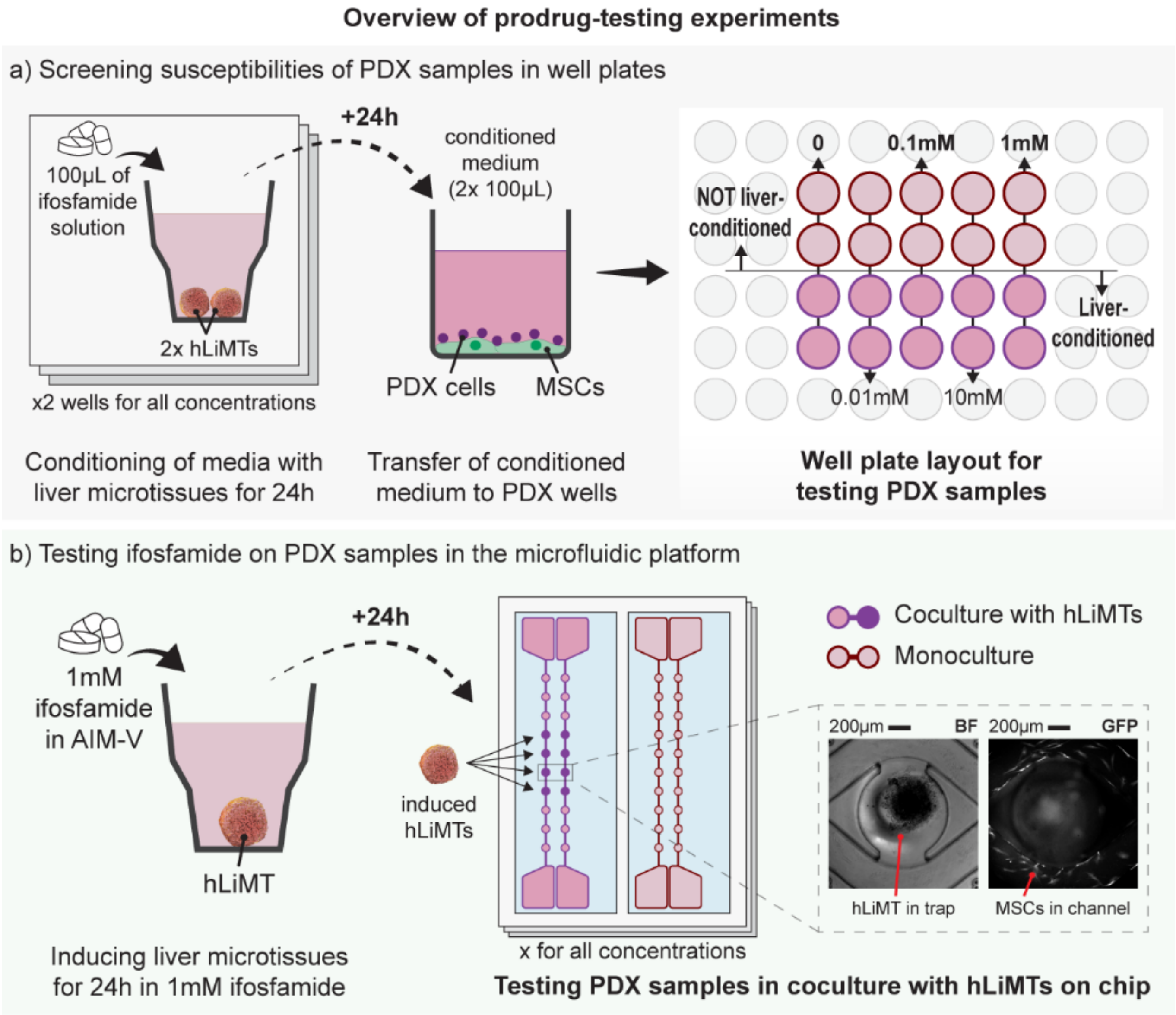


***Supplementary Figure 2.*** *Overview of prodrug testing experiments including hepatic bioactivation.* ***a)*** *Metabolic transformation of ifosfamide was mimicked in the traditional well-based assay by conditioning the drug solutions in two separate wells hosting two hLiMTs in 100 µL AIM-V each for 24 h and then transferring the conditioned medium to the PDX samples in the 96-well plate and incubation for 3 days.* ***b)*** *In order to include liver-metabolism effects in the microfluidic platform, the PDX samples were co-cultured with hLiMTs on chip (4 hLiMTs per channel) during the 3-day-long ifosfamide exposure. Microscope images show a bright-field (BF) image of a hLiMT in the elevated compartment (trap) and a GFP-channel image of the MSCs at the bottom of the interconnecting microfluidic channel of a prototype platform.*

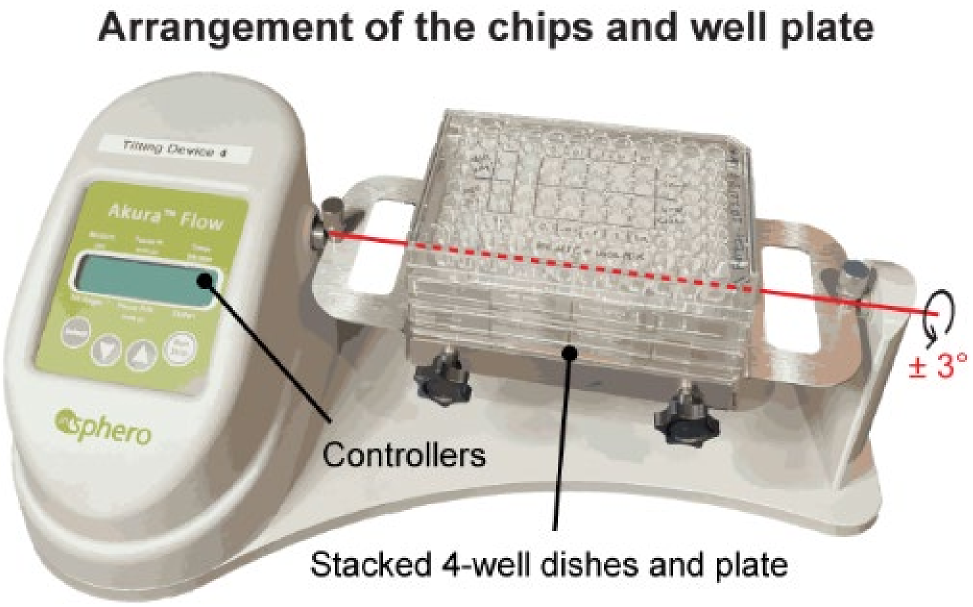


***Supplementary Figure 3.*** *Arrangement of the chips and well plates on a tilting device. During the 3-day-long ifosfamide treatment of PDX samples, plates accommodating 4 devices with replicates of the microfluidic platform and the 96-well plate for the traditional well-based testing were stacked on top of each other and tilted continuously in a cell culture incubator at 37°C, 95% humidity and 5%* $CO_2$.

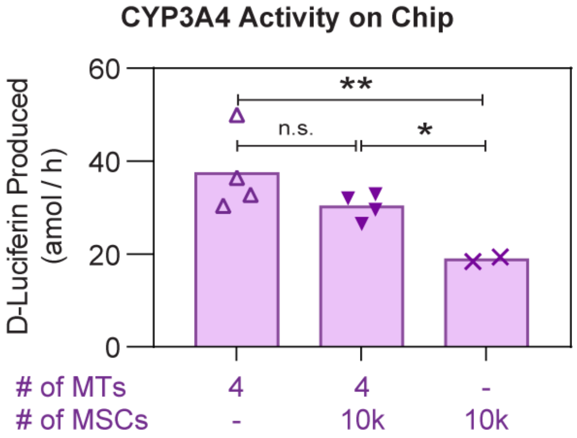


***Supplementary Figure 4.*** *Measured CYP3A4 activity in our platform using a P450-Glo CYP3A4 Assay. D-luciferin production per hour of four hLiMTs and 10k MSCs in monocultures or co-culture in our platform after 72 h (n=4 for 4-hLiMT monoculture and 4 hLiMTs and 10k MSCs in co-culture; n=2 for 10k-MSC monoculture). All hLiMTs for on-chip measurements were induced by culturing in 1 mM ifosfamide solution for 24 h before the CYP3A4 assay. Total D-luciferin production was normalized by 72 hours to estimate the average CYP3A4 activity throughout the experiment. * P<0.05, ** P<0.01*

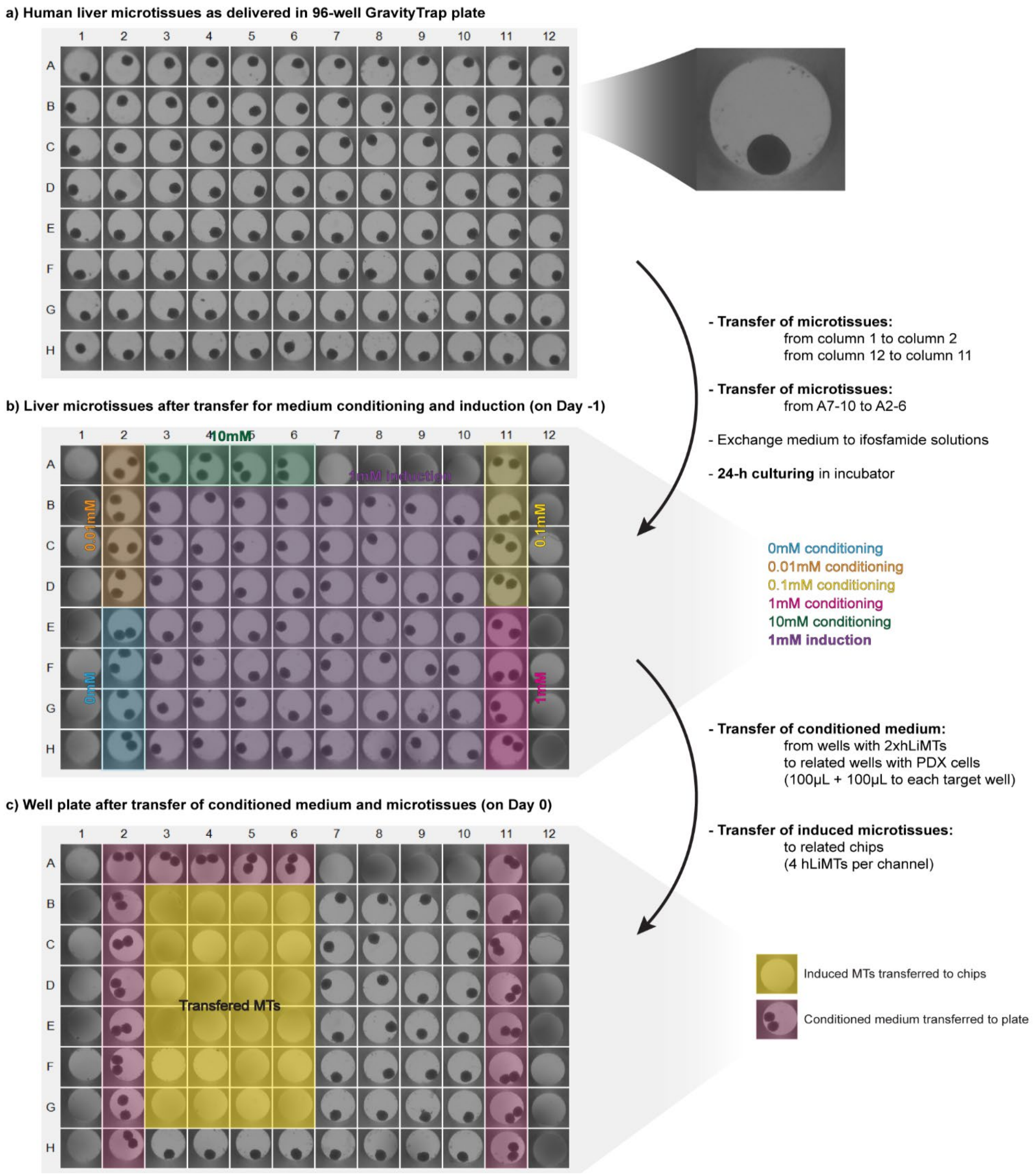


***Supplementary Figure 5.*** *Human liver microtissues (hLiMTs) during the experiment.* ***a)*** *The hLiMTs were delivered in a 96-well Akura96 well plate and were maintained in the hLiMM-AF liver maintenance medium until Day -1.* ***b)*** *On Day-1, hLiMTs in column 1 (A1 to H1), column 12 (A12 to H12), and in wells from A7 to A10 were transferred to the neighboring wells in column 2 (A2 to H2), column 11 (A11 to H11), and A3 to A6, respectively. The wells hosting 2 hLiMTs were used to condition the medium for well plate-based experiments. For each PDX sample that was treated on the well plate, 2 wells were hosting a total of 4 hLiMTs, each having 100 µL of the ifosfamide solution of the corresponding concentrations (0, 0.01, 0.1, 1 and 10 mM). The rest of the wells had 1 hLiMT each, with 1mM ifosfamide solution in AIM-V medium for inducing CYP3A4 activities.* ***c)*** *After 24h of incubation, on Day 0, the conditioned medium was transferred to the treatment wells with PDX samples (100 µL+100 µL to each target well) to*

*activate liver metabolism in the well-based protocol. The induced hLiMTs were transferred to the microfluidic chips that hosted PDX samples that were treated with ifosfamide for assessing the efficacy of ifosfamide and its metabolites.*

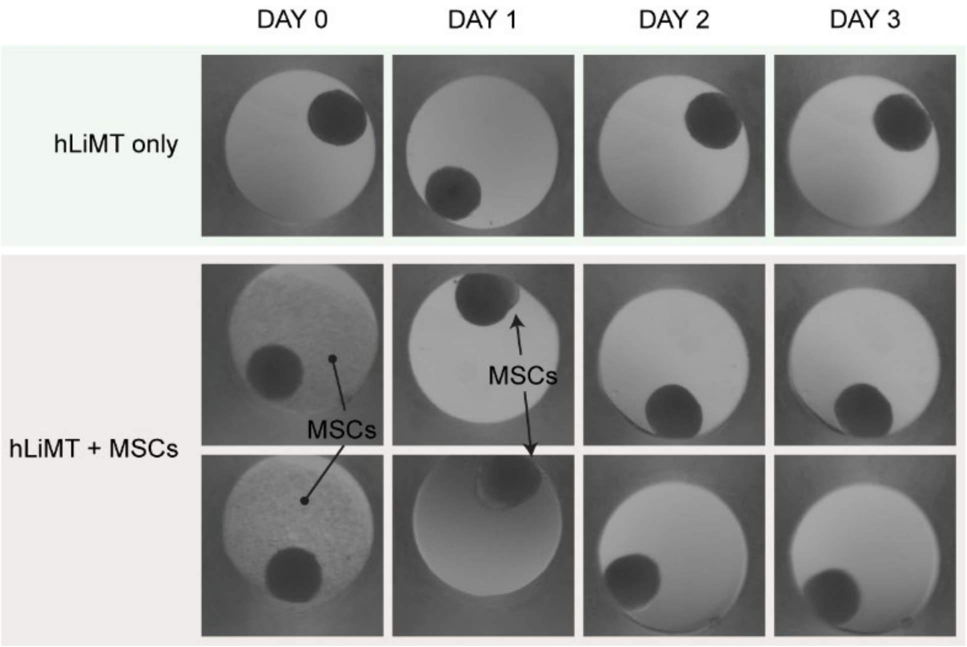


***Supplementary Figure 6** Co-culturing of MSCs and hLiMTs in a well plate for 3 days. TOP: hLiMTs in monoculture remained intact and maintained their shape. BOTTOM: When MSCs and hLiMTs were cultured together, MSCs first sedimented at the bottom of the wells (DAY 0) and later aggregated and merged with the hLiMTs. The aggregated MSCs can be seen especially on DAY 1 as they seemingly "deformed" the hLiMT shape by merging with the hLiMTs. On DAY 2 and DAY 3, the circular shape was recovered, as MSCs wrapped around the hLiMTs. Neither hLiMTs (hepatocytes) nor MSCs grew when a microtissue was formed.*

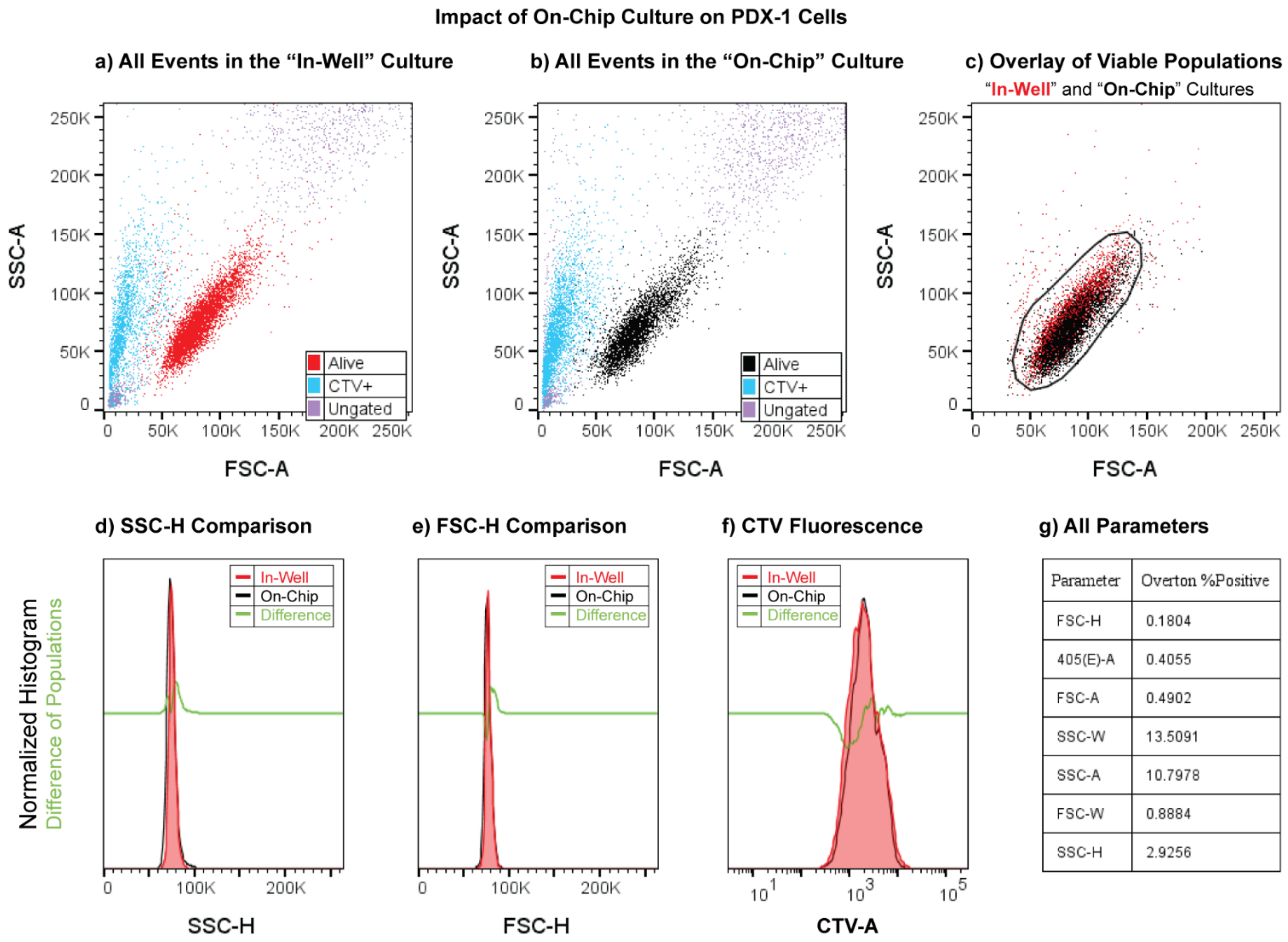


| Parameter | Overton %Positive |
|---|---|
| FSC-H | 0.1804 |
| 405(E)-A | 0.4055 |
| FSC-A | 0.4902 |
| SSC-W | 13.5091 |
| SSC-A | 10.7978 |
| FSC-W | 0.8884 |
| SSC-H | 2.9256 |

***Supplementary Figure 7.*** *Impact of on-chip culturing on PDX-1-cell morphology. All events were recorded with flow cytometry from* ***a)*** *"in-well" cultures and from* ***b)*** *"on-chip" cultures after 3 days of culturing without ifosfamide exposure. CTV+ cells (except viable PDX cells) are colored in* ***light blue****. Viable PDX-1 cells of "in-well" cultures are colored in* ***red****, whereas viable PDX-1 cells of "on-chip" cultures are colored in* ***black****.* ***c)*** *Overlay of the viable cell populations of the two culture platforms. The gating was based on the distribution of "in-well" cells and covered 97.3% and 96.7% of "in-well" and "on-chip" cell populations, respectively. Comparison of* ***d)*** *side scatter heights (****SSC-H****),* ***e)*** *forward scatter heights (****FSC-H****), and* ***f)*** *CellTrace Violet fluorescence amplitudes (****405(E)-A****) of the viable "in-well" and "on=chip" cell populations. Color coding of populations are the same as in the scatter plots (****red*** *for "in-well",* ***black*** *for "on-chip" populations, and* ***light green*** *for the difference).* ***g)*** *Statistical comparison of all relevant parameters in terms of Overton %Positive cells, where a lower percentage indicates higher similarity [40]. A coverage >95% was observed for the majority of investigated parameters, indicating no significant change in cell morphology due to "on-chip" culturing. Refer to* ***Supplementary Figure 13*** *for a detailed FACS gating strategy that was used to analyze the PDX populations.*

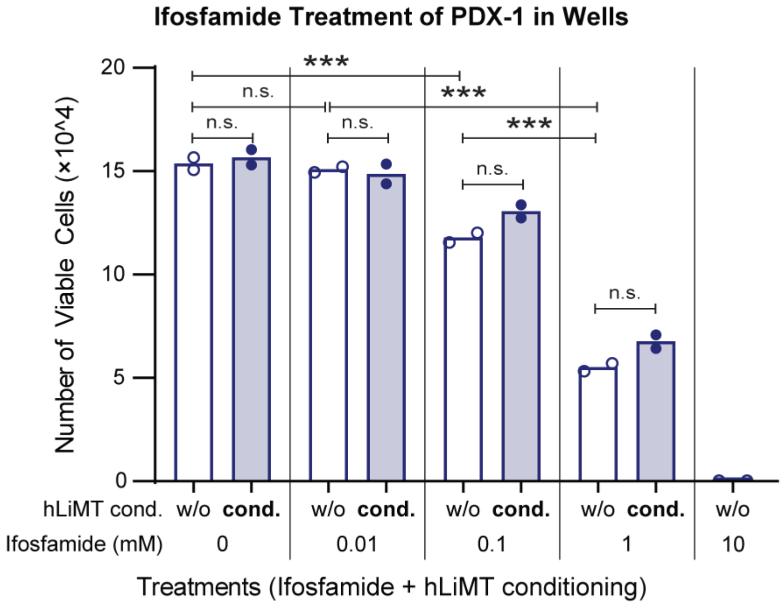


***Supplementary Figure 8.*** *Number of viable cells after ifosfamide treatment of PDX-1 cells in the well plate with or without hLiMT conditioning of the culture medium. The culture medium was conditioned by adding hLiMTs to the ifosfamide solutions in the wells as described in the Materials and Methods section. Conditioning with hLiMTs did not significantly affect the viability of PDX cells in comparison to no conditioning, regardless of the concentration of ifosfamide. The number of viable cells decreased with increasing ifosfamide concentration, except for 0.01 mM ifosfamide that showed no significant impact on viability. Exposure to 10 mM ifosfamide resulted in the death of all cells. The number of cells (plotted biological replicates) was corrected, as the total number of viable cells has been obtained by using fluorescent counting beads; each data point represents the mean of 2 technical replicates of two distinct biological replicates of each treatment. *** P<0.001, analyzed using nested 1-way ANOVA.*

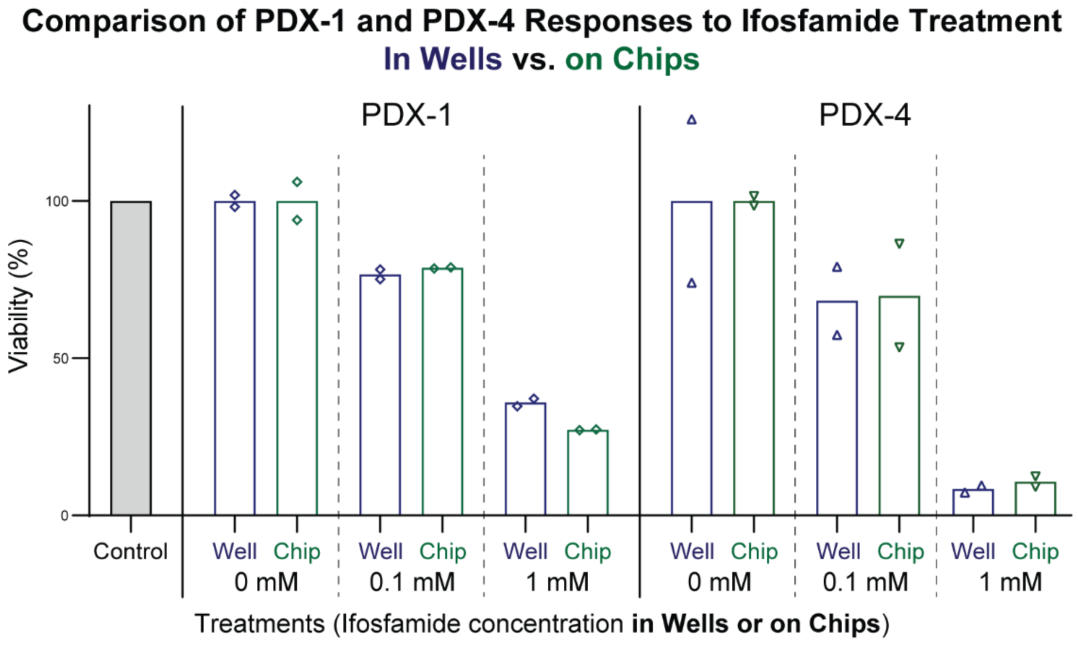


***Supplementary Figure 9.*** *Comparison of well-plate and on-chip ifosfamide responses of PDX-1 and PDX-4 samples* ***in the absence of hepatic metabolism*** *(as percentage of the control conditions for each PDX sample). For both samples, the on-chip responses were in agreement with the responses observed in the traditional well-based assay. This finding suggests that the microfluidic platform can be used to screen drug responses of patient-derived samples, similar to the well-based assays. It also shows that culturing PDX samples in the microfluidic platform during the screening does not alter the outcome of experiments.*

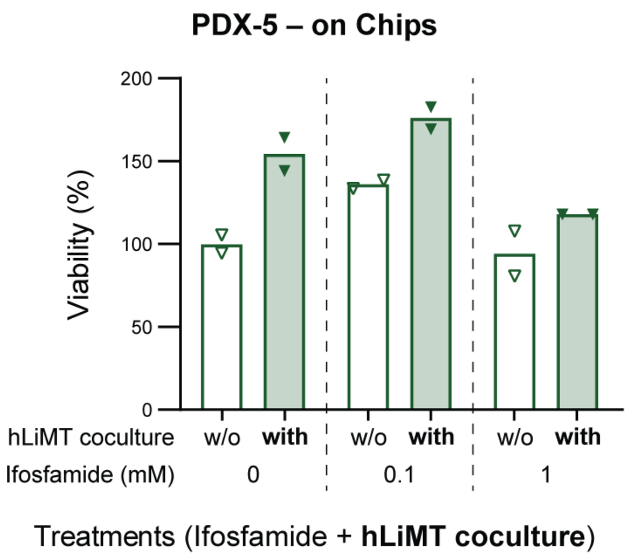


***Supplementary Figure 10.*** *Ifosfamide treatment of PDX-5 in our platform (as a percent of control condition –the treatment with 0 mM and no hLiMT in co-culture). Because of low on-chip viability, the response could not be profiled, and this sample was excluded from the analyses. Each data point in the bar plots represents the mean of two technical replicates (n=2) of experimental repetitions of each treatment in wells or on chips (n=2).*

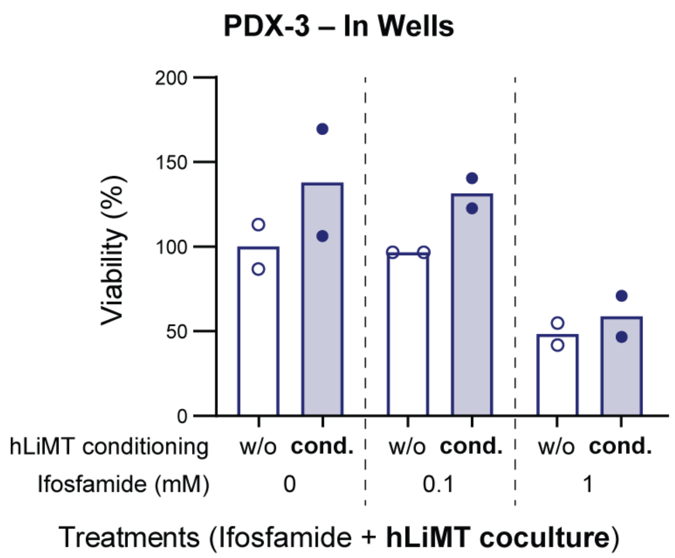


***Supplementary Figure 11.*** *Ifosfamide treatment of PDX-3 samples in the well plate (as percentage of the control condition –the treatment with 0 mM and no hLiMT in co-culture). Due to the low in-vitro viability, the response could not be profiled. Each data point in the bar plot represents the mean of two technical replicates (n=2) of experimental repetitions of each treatment in wells or on chips (n=2).*

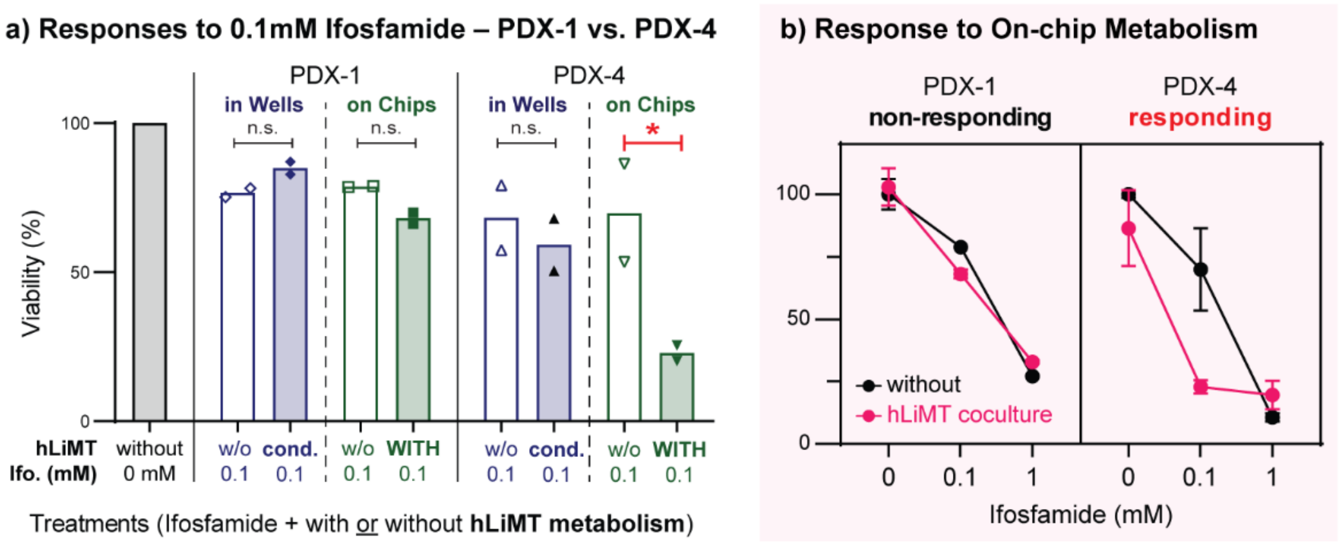


***Supplementary Figure 12. a)*** *Comparison of PDX-1 and PDX-4 viability after exposure to 0.1 mM ifosfamide in the well plate and in the microfluidic platform in the presence or absence of hepatic metabolism. For both PDX samples, the viability was similar with or without liver conditioning (with respect to the respective controls) in the well plate-based experiments. For PDX-1, on-chip co-culturing with hLiMTs did not result in a decreased viability. In contrast, the viability of the PDX-4 in on-chip co-culture with hLiMTs (i.e., with continuous metabolic transformation of ifosfamide) decreased significantly in comparison to the same treatment without hLiMT co-culturing. The PDX-4 response indicated that the prodrug was metabolized by the liver, which suggests that the microfluidic platform can be used to test the efficacy of prodrugs with patient-derived samples.* ***b)*** *Comparison of on-chip dose-responses of PDX-1 (non-responding to metabolites) and PDX-4 (responding) in our platform, when the samples were cultured with and without hLiMTs for hepatic activation. Each data point in the bar plots represents the mean of two technical replicates (n=2) of experimental repetitions of each treatment in wells or on chips (n=2). Each marker in the dose-response curve represents the mean of the data points of the bar plots, and error bars indicate the SEM (n=2). * P<0.05.*

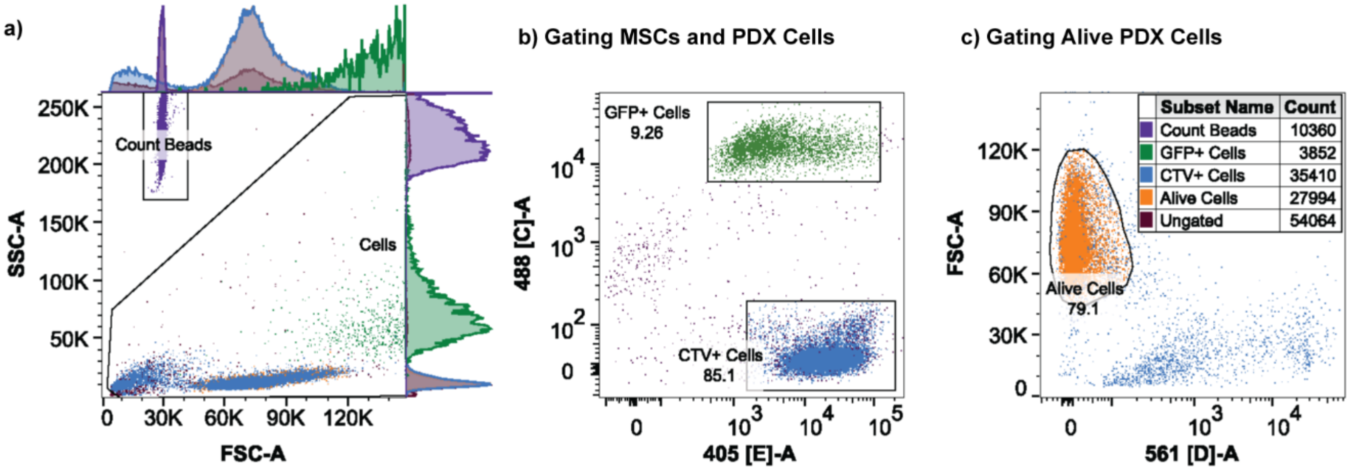


***Supplementary Figure 13.*** *FACS analysis and used gating strategy.* ***a)*** *Scatter plot of all recorded events, overlaid with populations (side scattering amplitude vs. forward scattering amplitude): counting beads (violet), CellTrace-Violet-positive cells (CTV-positive cells), and green-fluorescent-protein-expressing cells (GFP-positive cells).* ***b)*** *MSCs and PDX cells were gated with respect to their GFP expression (488 nm laser) or CTV amplitudes (405 nm laser). GFP-positive cells were identified as MSCs, and CTV-positive cells were identified as PDX cells. The counting beads were not included at this step of the analysis.* ***c)*** *Identification of alive PDX cells according to propidium-iodide signal amplitudes (561 nm laser). PI-negative cells were counted as viable cells and were used as a basis for the analysis. INSET: The legend for all shown populations.*

***Supplementary Table 1*** *Characteristics of samples included in this study. F: female, M: male, HR: high risk, SR: standard risk. The number of initially seeded viable cells at day 0 is based on hemocytometer/automated cell counting with DAPI staining. The counts at day 3 are averages of replicates under control conditions in wells and on chips (0 mM, without hLiMT co-culture). The number of seeded cells in wells and on chips were the same for PDX-1 and PDX-4. For PDX-2 and PDX-3 treatments in the well plate, the number of viable cells in the wells were normalized to the ratio of the initially seeded number of cells in the wells and that on the chips to compensate for slight differences in cell numbers.*

| Sample | Sex | Age | Risk stratification BFM | Number of viable cells | | |
|---|---|---|---|---|---|---|
| | | | | At Day 0 (same number of cells seeded for wells & chips) | At Day 3 | |
| | | | | | In wells | On chips |
| **PDX-1** | M | 6.5 | HR | ~100k | ~150k | ~82k |
| **PDX-2** | M | 8 | SR | ~116k | ~132k | ~37k |
| **PDX-3** | M | 14.3 | HR | ~62k | *LOW - excluded* | ~1.8k |
| **PDX-4** | M | 2 | SR | ~100k | ~22k | ~9.5k |
| **PDX-5** | F | 11.2 | HR | ~80k | *No data* | *LOW - excluded* |

***Supplementary Table 2*** *Summary of contrasts as discussed in the manuscript and statistical analyses for the relevant treatments.*

| Sample | Platform | hLiMT | IF | Contrast | Significance p-value | | *df* | 95%-CI (Lower CL – Upper CL) | |
|---|---|---|---|---|---|---|---|---|---|
| **PDX-1** | Well | - | 0 | hLiMT: w/o – with | n.s | P>0.05 | 12 | -0.199 | 0.160 |
| | Chip | - | 0 | hLiMT: w/o – with | n.s. | P>0.05 | 12 | -0.208 | 0.152 |
| | Well | w/o | - | IF: 0.1 – 0 | ** | 2.806e-03 | 12 | -0.445 | -0.086 |
| | Well | w/o | - | IF: 1 – 0.1 | *** | 2.881e-09 | 12 | -0.938 | -0.578 |
| | Well | - | 0.1 | hLiMT: w/o – with | n.s. | P>0.05 | 12 | -0.282 | 0.078 |
| | Chip | - | 0.1 | hLiMT: w/o – with | n.s. | P>0.05 | 12 | -0.035 | 0.325 |
| | - | w/o | 0.1 | Platform: Well – Chip | n.s. | P>0.05 | 12 | -0.206 | 0.153 |
| **PDX-4** | Well | - | 0 | hLiMT: w/o – with | n.s | P>0.05 | 12 | -0.814 | 1.209 |
| | Chip | - | 0 | hLiMT: w/o – with | n.s. | P>0.05 | 12 | -0.851 | 1.173 |
| | Well | w/o | - | IF: 0.1 – 0 | n.s. | P>0.05 | 12 | -1.372 | 0.652 |
| | Well | w/o | - | IF: 1 – 0.1 | *** | 1.429e-04 | 12 | -3.098 | -1.074 |
| | Well | - | 0.1 | hLiMT: w/o – with | n.s. | P>0.05 | 12 | -0.872 | 1.152 |
| | Chip | - | 0.1 | hLiMT: w/o – with | * | 3.050e-02 | 12 | 0.082 | 2.105 |
| | - | w/o | 0.1 | Platform: Well – Chip | n.s. | P>0.05 | 12 | -1.019 | 1.004 |
| **PDX-2** | Well | - | 0 | hLiMT: w/o – with | n.s | P>0.05 | 12 | -0.258 | 0.709 |
| | Well | - | 0.1 | hLiMT: w/o – with | n.s. | P>0.05 | 12 | -0.453 | 0.514 |
| | Chip | - | 0 | hLiMT: w/o – with | n.s. | P>0.05 | 12 | -0.325 | 0.641 |
| | Chip | - | 0.1 | hLiMT: w/o – with | n.s. | P>0.05 | 12 | -0.439 | 0.527 |
| **PDX-3** | Chip | - | 0 | hLiMT: w/o – with | n.s. | P>0.05 | 6 | -1.522 | 1.895 |
| | Chip | - | 0.1 | hLiMT: w/o – with | n.s. | P>0.05 | 6 | -1.617 | 1.799 |

*IF: Ifosfamide, CI: Confidence Interval, CL: Confidence Level, with (w/o): with (without) metabolism, df: degrees of freedom*
